# Regionalization of gene expression and cell types in the silk gland of the pantry moth *Plodia interpunctella*

**DOI:** 10.1101/2025.07.11.664249

**Authors:** Jasmine D. Alqassar, Mathilde Biot, Lauren E. Eccles, Whitney L. Stoppel, Arnaud Martin

**Author notes:** Corresponding authors : Jasmine Alqassar,; Arnaud Martin.

## Abstract

Lepidopteran silk glands include a posterior silk gland (PSG) that secretes core fibers and a middle silk gland (MSG) that secretes adhesive sericins. While well-studied in the silkworm (*Bombyx mori*), little is known about the gene expression profiles underlying the diversity of lepidopteran silks. Here we characterized the silk gland from the pantry moth *Plodia interpunctella* using a combination of quantitative and spatial assays. RNA-seq analyses of differential gene expression between the MSG and PSG depict the transcriptomic divergence between these two secretory tissues. In the PSG, high expression of fibroin genes *FibL* and *FibH* were detected, whereas MSG samples were dominated by transcripts encoding major sericins— bioadhesive proteins that form the coating of the silk fiber. Hybridization Chain Reaction (HCR) mRNA profiling revealed sharp cellular boundaries within the silk gland: PSG cells exclusively expressed fibroins, while MSG comprised two compartments each expressing different combinations of sericins. Our findings corroborate the conserved organization of lepidopteran silk glands into specialized secretory subdivisions, and establish *Plodia* as a promising comparative model for studying silk diversity. This work provides a foundation for future research into the cellular and evolutionary basis of silk production across Lepidoptera.

## Introduction

There are an estimated 180,000 species of moths and butterflies (Lepidoptera) and most produce silk throughout their larval life, with common functions including locomotion, protection, habitat construction, and pupation. Silk is a complex biomaterial formed through protein self-assembly in a pair of labial glands (the silk glands), where silk proteins are secreted and then extruded through the spinneret near the mouth^1,2^. In Lepidoptera, silk fibers are characterized by a fibroin core with an outer protein coating^3–5^. The inner fibroin core provides the primary structural strength and toughness of the silk fiber^6–9^, while the outer coating modulates properties such as hydrophilicity and adhesion^5,10–13^. Each lepidopteran organism that produces silk fibers regulates the composition of these layers through the expression of secreted proteins in their silk glands^4^.

Investigations in *Bombyx mori* have outlined the general structure of the silk gland as divided into three distinct regions, each with specialized functions related to silk production^5,14,15^. The posterior silk gland (PSG) is primarily responsible for the production and accretion of proteins that form the core of the silk fiber: Fibroin Heavy chain (FibH), Fibroin Light chain (FibL), and Fibrohexamerin (Fhx, also known as P25). FibH contains repetitive motifs that create crystalline domains and underlie the fiber’s mechanical strength^1,9,16–20^. The C-terminal region of FibH forms covalent bonds with FibL, which acts as an essential linker to form silk crystalline units^21,22^. Fhx is dispensable for silk production but is thought to assist with the trafficking of the fibroin proteins before they accumulate within the lumen of the gland^23,24^. The middle silk gland (MSG) is primarily responsible for the synthesis of sericin proteins, a family of hydrophilic adhesive proteins that form a coating layer around the fibroin core^11^. In the silkworm *Bombyx mori*, gland morphology and sericin expression profiles of the silkworm MSG suggest it is itself subdivided into three subregions called the anterior, middle, and posterior MSGs, abbreviated MSG-A, MSG-M, and MSG-P^25–29^. Last, the anterior silk gland (ASG) consists in a thin duct involved in silk fiber alignment and stretching, before extrusion through the spinneret^25,26,30^. The ASG is also connected to a small pair of exocrine structures called Filippi’s glands^31^, and this accessory gland likely contributes to silk maturation and cocoon compaction in *Bombyx*^32^.

While the silk of the domesticated silkworm *Bombyx mori* has been extensively characterized, it represents only a single lineage within the vast diversity of Lepidoptera, and expanding this research to more species hold great promise to uncover the proteins that underlie the mechanical properties^33^, adhesivity^24^, and antimicrobial activity of silks specialized to different ecological uses^34–36^. This comparative endeavor has been spurred by the generation of silk gland transcriptomes in an increasing number of lepidopteran lineages^37–42^. However, few of these species benefit from state-of-the-art gene annotation resources, and because spatial assays that directly visualize gene products in dissected glands have been limited to *Bombyx*^12,16,27–32^, these studies often offer a limited resolution on how gene expression varies in the silk gland.

The pantry moth, *Plodia interpunctella*, is a promising alternative model system for the study of silk biology, as recent work has highlighted its potential in silk fiber production for biotechnology applications^49–51^, as well as a laboratory powerhouse for genetic manipulation^52–55^. To expand the understanding of the structure and organization of the *Plodia* silk gland, we carried out an RNAseq differential gene expression analysis of the MSG and PSG. We selected this species as a model system for silk biology, due to the rearability of this insect, its potential for functional genomics^53–55^, and the availability of an annotated genome. As a pyralid moth, *Plodia* is also phylogenetically close to *Ephestia kuehniella* and *Galleria mellonella*, where comparative studies of gland histology and gene expression suggest the MSG subdivides into two domains of sericin expression^10,31^. To test this in *Plodia*, we leveraged the spatial specificity of Hybridization Chain Reaction (HCR), a technique that amplifies the fluorescent signal of small probes targeted at an mRNA of interest, and refined the expression patterns of silk factor genes in whole-mount glands with a subcellular resolution. We discuss how combining RNAseq and HCR primes future explorations of silk diversity in Lepidoptera and beyond.

## Results

### Morphology and cellular organization of the *Plodia* silk glands

We used dissections and high-resolution micro-computed tomography (micro-CT) of fifth instar larvae to reveal the positioning and overall morphology of the *Plodia* silk glands **(Fig. 1A**-**1C)**. Each gland forms an epithelial tube connected to the mouth spinneret via a thin ASG, followed by a MSG thick section resting ventrally to the gut and starting in the T2 mesothoracic segment. The two glands curve backwards when reaching the A4 abdominal segments, and then curve back towards the posterior end in the A2 segment. This second curve approximately marks the beginning of the two PSGs, which run in a position lateral to the gut from the A3 to A5 segments.

**Figure 1.**
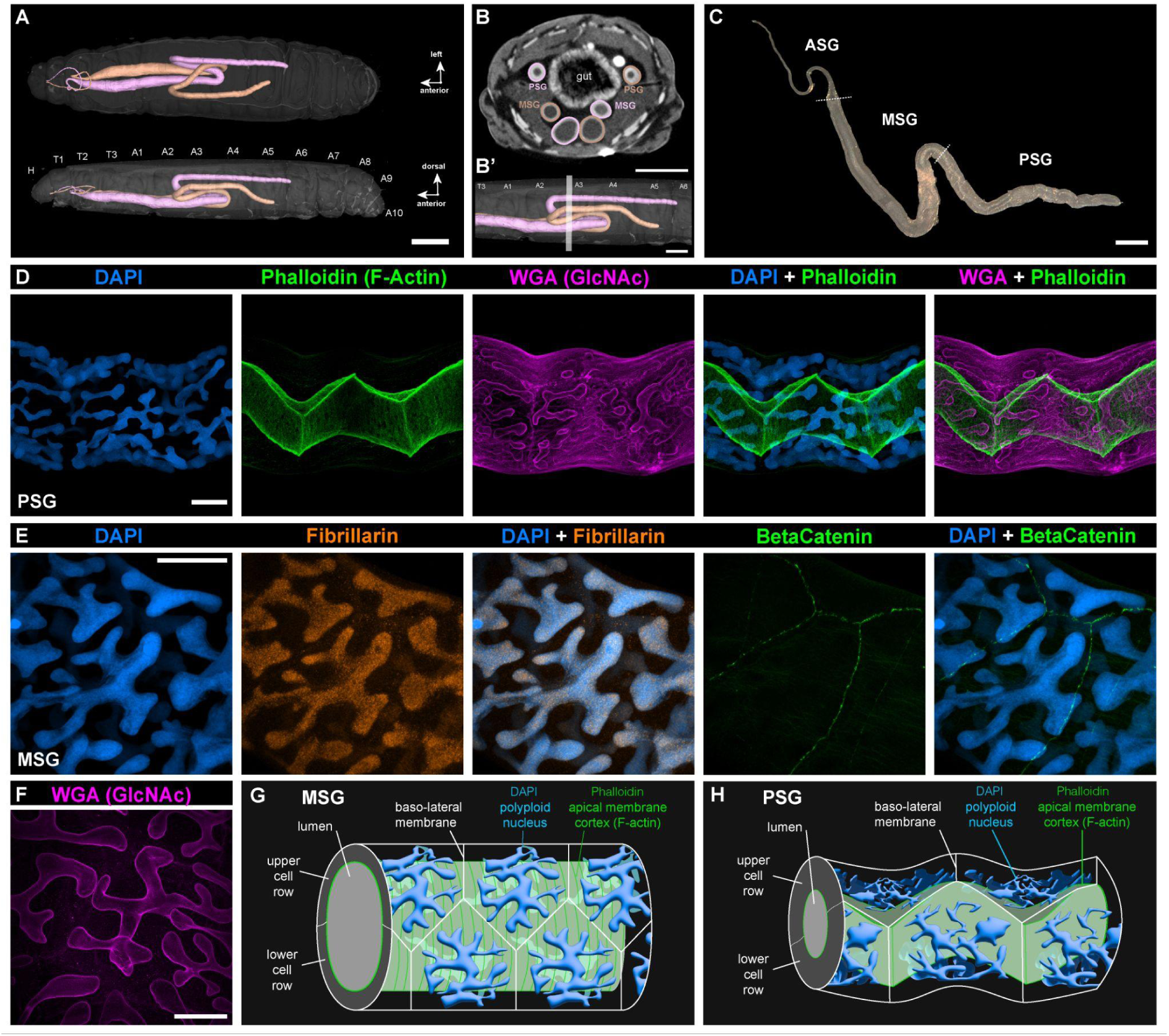
Cellular morphology of *Plodia* silk glands. **A.** Micro-CT scan of a fifth instar larva with each silk gland rendered in 3D. False coloring marks the identity of each individual glands. Glands switch sides twice in this individual (in T2 and A3), but this crossing does not always occur. Top : ventral view ; bottom : lateral view. Animated version in **Video S1**. **B-B’.** Cross-sectional view of a scanned larva in the A3 segment (section plane as shown in B’), where the PSG and two sections of the MSG are visible. Gland outer and inner diameters both increase as the gland progresses towards the ASG. Animated version in **Video S2. C.** Dark-field image of a dissected silk gland from a *Plodia* fifth instar wandering larva. Dotted lines indicate the ASG/MSG (top) and MSG/PSG (bottom) boundaries. **D.** DAPI, Phalloidin (F-Actin), and WGA (GlcNAc) staining of a wandering fifth instar PSG. Comparisons with ASG and MSG sections are featured in **Figure S1** and **Video S3**. **E.** DAPI and monoclonal antibody staining for Fibrillarin (marking nucleoli), and Beta-Catenin (accumulating at apical cell membrane junctions) in a fifth instar *Plodia* MSG. **F.** WGA staining of the nuclear envelope in polyploid nuclei of the MSG. **G-H.** Schematic representation of cell organisation and geometry in the MSG (G), and PSG (H). Scale bars : A = 1 mm; B-B’, C = 500 μm ; D-F = 50 μm.

Next we visualized the cellular morphology of the PSG and MSG using confocal microscopy **(Fig. 1D; Fig. S1)**, using DAPI stainings of nuclear DNA, phalloidin stainings of F-Actin, and WGA stainings of N-acetylglucosamine (GlcNAc, labeling various membranal features). Phalloidin marked the apical surface of all cells along the silk gland, thus effectively contouring the luminal space left by cells. Packing of the single-cell layer results in 120° angles at the junctions between three cells, characteristic of epithelial tubes^56^, in both the PSG and MSG. Apical cell surfaces show an accordion shape in the PSG, likely due to the small size of the lumen and the relatively small diameter of the gland **(Fig. 1D; Fig. S1C; Video S3).** In the MSG, as the lumen has increased in diameter **(Fig. S1B),** and the cell apical surfaces that line it up are stretched around its periphery **(Fig. S1B; Video S3)**. Apical F-actin formed transverse striations forming ring-like structures around the lumen (perpendicular to the gland longitudinal axis). This overall geometry is consistent with the previously identified cellular morphology of the ASG, MSG, and PSG in silkworms^57^.

DAPI stainings show that MSG and PSG nuclei are extremely large and elongated (**Fig. 1D-E**), with a branching architecture also visible in WGA stainings of the nuclear envelope^58^. Such large nuclei are typical of silk glands across Lepidoptera and are caused by endoreplication, a mode of cell cycling characterized by genome replication leading to an increase in cell size without cell or nuclear division^59–62^. The branching architecture increases the surface-to-volume ratio of nuclei, and was proposed to optimize the efficiency of gene expression and transcript processing in these large cells^62^. Stainings of silk gland nucleoli using an anti-Fibrillarin antibody show a constellation of signal puncta **(Fig 1E)**, contrasting with the 1-4 nucleoli expected in diploid cells. While we could not quantify these signals to measure ploidy levels reliably, a previous study of the equivalent cells in *Ephestia* last-instar larvae measured that silk gland cells accumulated over 8,000 genome copies^62^. Cells and nuclei are prone to bursting due to their enormous size, and we were unable to isolate intact silk gland cells or live nuclei using cell-dissociation methods optimized for less polyploid (4n-32n) cells from developing pupal wings^63,64^, which precluded us from attempting single-cell transcriptomics in this project.

### Hybridization Chain Reaction reveals a sharp boundary between MSG and PSG

We sought to test the suitability of HCR RNA *in situ* hybridization^65^ to detect gene expression at subcellular resolution within the *Plodia* silk gland, initially targeting the intronic and exonic sequences of *Fibroin Heavy Chain* (*FibH*) (**Table S1**). Both probe sets showed a strong signal in PSG cells: the intronic probe signal was restricted to the nucleus, consistent with the detection of nascent transcripts, while exonic probes showed a diffuse signal of the mRNA in the nucleus and cytoplasm (**Fig. 2A**). *FibH* intronic and exonic stainings, as well as *FibL* (*Fibroin Light Chain*) mRNA staining, were exclusive to the PSG and revealed a strong spatial boundary with the adjacent MSG (**Fig. 2B-C**). *FibH* and *FibL* mRNA expression colocalizes in the PSG and have prominent boundaries of expression. Unexpectedly, staining for *P25/Fibrohexamerin* (*Fhx*) showed prominent expression in the MSG and a weaker signal in the PSG (**Fig. 2D-E**). This result contrasts with silkworms, where *Fhx* is restricted to the PSG^44,66^, and where it is dispensable for silk fibroin assembly but participates in its secretion^23^. Fhx may fulfill a secretory function in both the MSG and PSG in *Plodia*, and is lacking in the ASG, thereby delineating a sharp ASG/MSG boundary between non-exocrine and exocrine sections of the silk gland (**Fig. 2D**). Overall, these results posit HCR as a powerful spatial assay for profiling gene expression in whole-mount silk glands, and highlight sharp functional boundaries in the silk gland.

**Figure 2.**
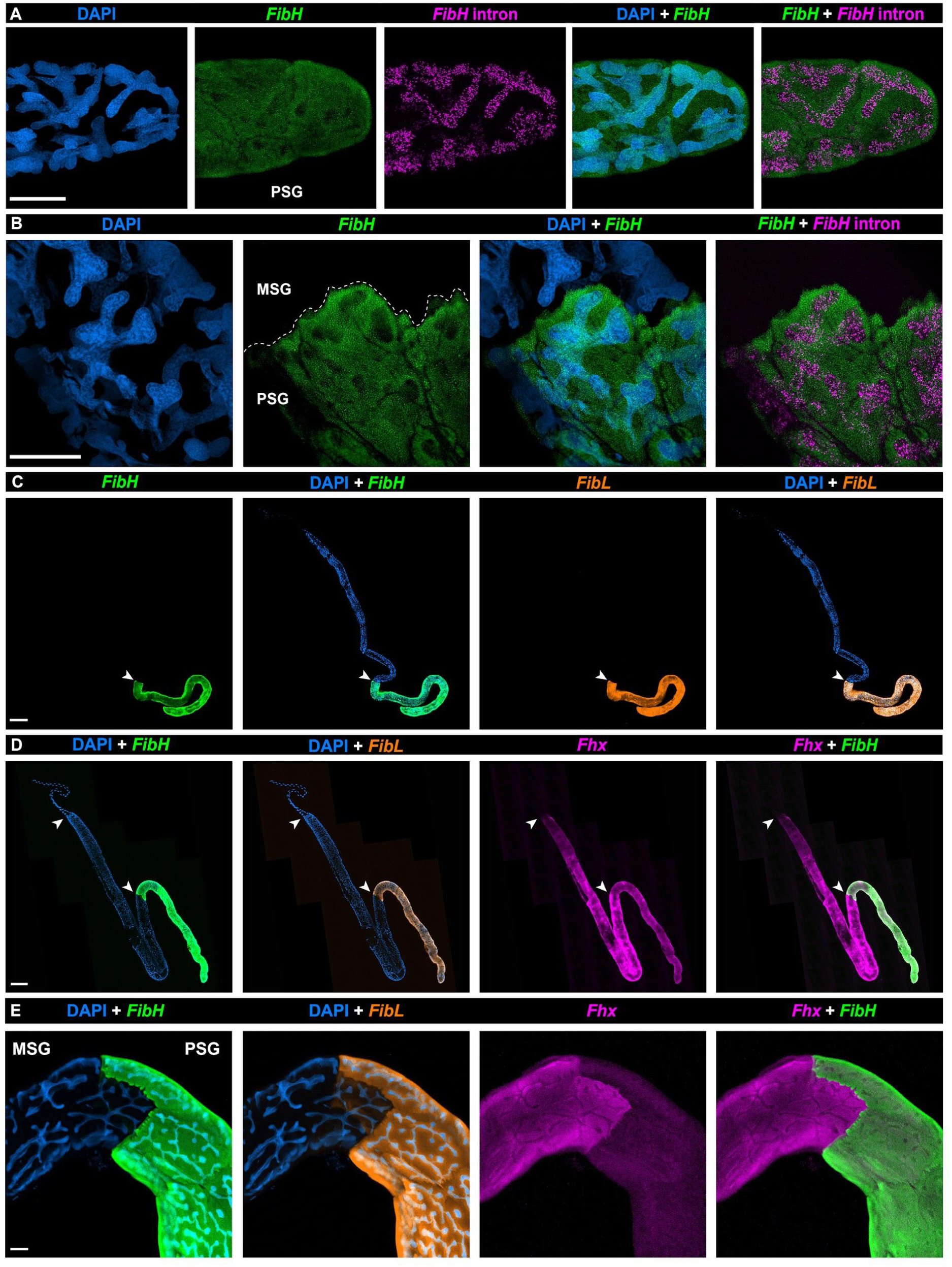
HCR detection of fibroin transcripts reveals tissue boundaries in the silk gland. **A-B.** HCR stainings using exonic and intronic *FibH* probes in the posterior PSG (A) and at the MSG/PSG interface (B). **C.** Whole-mount fifth instar silk gland HCR stainings for *FibH* and *FibL*. **C-E.** HCR stainings of wandering fifth instar silk glands for *FibH, FibL* and *Fhx* mRNA. Arrowheads: MSG/PSG (C-D: bottom) and ASG/MSG boundaries (D: top). Scale bars : A-B, E = 50 μm.; C, D = 500 μm.

### RNAseq of the PSG and MSG uncovers the major silk factors of *Plodia* silk

Next, we generated deep transcriptomes of the MSG and PSG of fifth instar larvae, as well as salivary glands and whole larval heads from the same individuals (four biological replicates per tissue, 16 libraries total). Dissected silk glands were manually split at the approximative MSG/PSG boundary using fine dissecting scissors before RNAseq, resulting in MSG samples (including portions of the ASG, but not the Filippi’s Glands), and PSG samples. In order to increase the statistical power of differential expression analyses, salivary gland samples were included as an exocrine tissue not involved in silk production, and larval heads as a control encompassing a heterogeneous mixture of cell types. DESeq2 analysis detected a total of 2,914 Differentially Expressed Genes (DEGs) between the MSG and PSG (**Table S2** ; adjusted *p* < 0.05). Out of the 229 top DEGs — defined here as those with the highest expression differences (log_2_FC > 3) — 23 were specific to the PSG (**Fig. 3A; Table S2-S4**), including *FibH*, *FibL,* and *Arrowhead* (*Awh*), a transcription factor known to regulate fibroin genes in silkworms^67,68^. Out of 229 top DEGs, 195 were enriched in the MSG relative to the PSG (**Fig. 3A; Table S2-S4**). Among them, genes that have been associated with the MSG of pyralid moths such as *Ser3a*, *Mucin12*, *SerP150*, *MG4, and Zon1*^10,12,69^, as well as the homeobox gene *Antp* known to determine the MSG identity in silkworms^28,29,44^, were restricted to the MSG.

**Figure 3.**
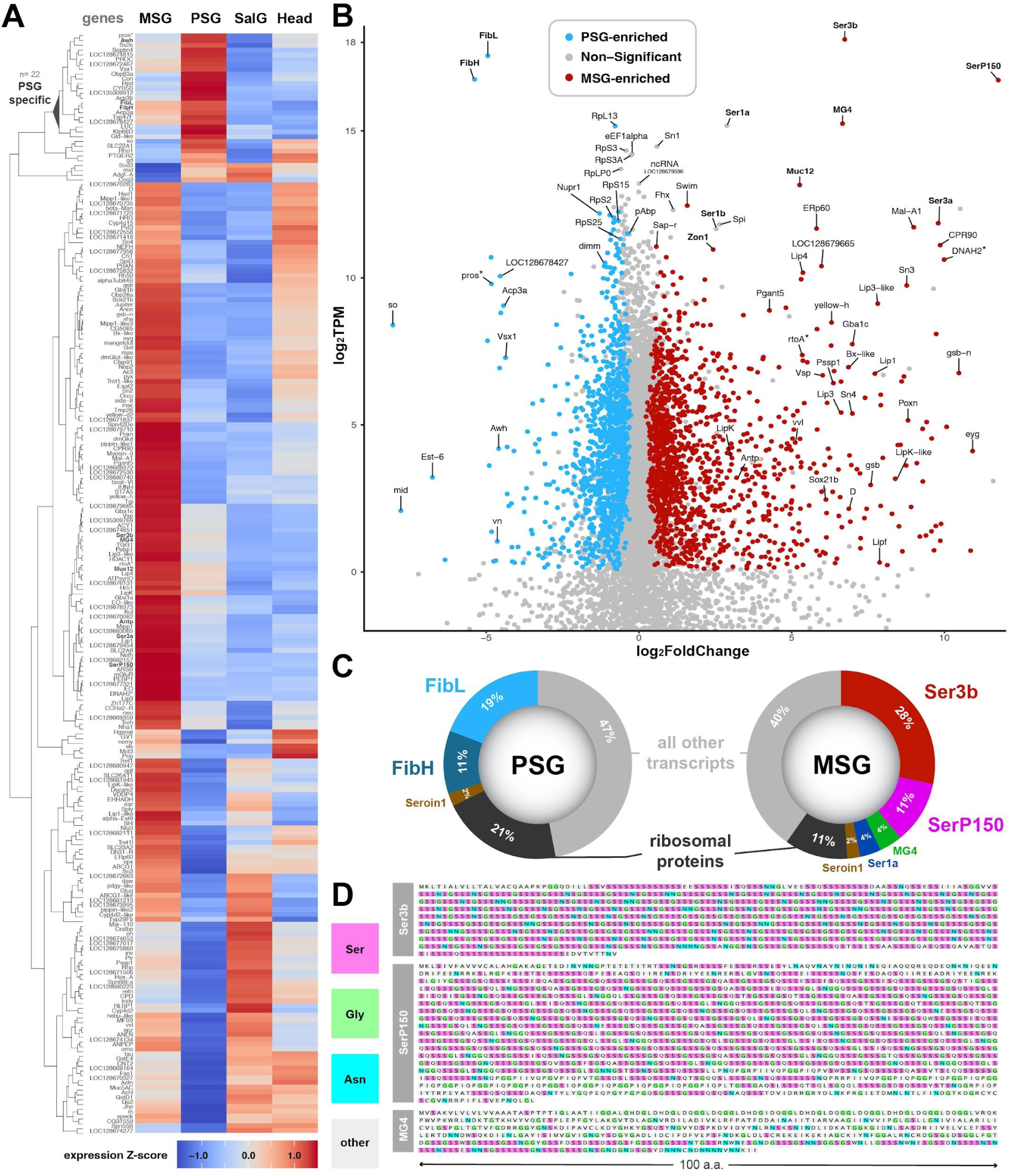
RNAseq profiling of transcriptome divergence in the PSG and MSG specialized glands. **A.** Heatmap of the top 229 DEGs between MSG and PSG (adjusted *p* < 0.01, |log_2_FoldChange| > 3, DESeq2 normalized counts > 300). Gene expression profiles across tissues are shown with a Z-score transformation and sorted by hierarchical clustering (SalG: salivary glands ; Heads: larval heads). **B.** Scatter plot highlighting expression level difference of individual genes between the PSG and MSG (x-axis, log2FC), and their relative transcript abundance within their tissue of enrichment (y-axis, log_2_TPM). Coloring highlights genes with DESeq2 adjusted *p* < 0.05 and log_2_TPM > 0.01. Non-significant, lowly-expressed genes (log_2_TPM < -2) are not shown. Asterisks: the genes *prospero* (*pros*), *rtoA*, and *DNAH2* are respectively adjacent to DEGs *FibH, MG4*, and *SerP150*, which may drive their enrichment in the corresponding tissues. **C.** Transcript representation within the PSG and MSG tissues (TPM as %). Ribosomal protein genes were pooled. **D.** Amino-acid composition of three major secreted proteins detected in the MSG, each showing extensive serine-rich stretches and repeats characteristic of sericin proteins. Accession numbers: XP_053622673 (Pi_Ser3b), XP_053622719 (Pi_MG4), XP_053613126 (Pi_SerP150).

We hypothesized that secreted silk factors are expressed at higher levels than non-secreted products. In addition to fold-change expression differences between the MSG and PSG, we used mean TPM (transcripts per million reads) values as a measure of transcript abundance within tissues^70^ (**Fig. 3A; Table S5**). A scatter plot summarizes this combination of relative expression within tissues and fold-change expression differences between tissues (**Fig. 3B; Table S6**). Both transcriptomes were strongly biased toward a small set of highly expressed secreted factors (**Fig. 3C**): *FibH* and *FibL* are by far the two highest expressed genes in the PSG transcriptome, amounting to 30% of all transcripts molecules sequenced in this tissue, and four sericin genes named *Ser3b*, *SerP150, MG4* and *Ser1a* (**Fig. 3D, Figs. S2-S5; Table S7**) represent 49% of the transcript molecules sequenced in the MSG. *Ser3b* alone accounts for 28% of all transcript molecules detected in the MSG, making this paralog of *Ser3a* (**Fig. S5**) a major component of the *Plodia* silk coating layer. Three other sericin genes (*Ser1b*, *Ser3a* and *Mucin12*) were also detected in the MSG at lower levels (0.3-0.9%).

Both PSG and MSG tissues showed high expression of ribosomal proteins, which together constitute 21% and 11% of the PSG and MSG transcriptomes, as well as of the protein elongation factors *eEF1alpha1*, *eEF1beta*, *eEF1gamma*, *eEF2*, *eEF5* and the *pAbp* translation termination factor, each with a log_2_TPM > 10 (**Fig. 3C**, **Table S5-S6**). Of note, both the PSG and MSG also expressed the seroin gene *Seroin1* (*Sn1*, log_2_TPM > 14.4) as well as a silk gland-specific non-coding RNA of unknown function (*LOC128679596*, log_2_TPM > 12.7). With the exception of their main fibroin and secreted factors, the most highly expressed transcripts were thus shared between the MSG and PSG.

### Candidate determinants of ASG/MSG/PSG specialization at the transcriptional and post-translational levels

Candidate genes for the regulation of transcription showed marked divergence between the PSG and MSG, some of which likely underlie the partitioning of the silk gland into at least two distinct exocrine tissues. Indeed, several transcription factors (TFs) were differentially expressed between the two regions, and may be important regulators of their identity (**Fig. 3B)**. In the PSG, this included genes encoding Awh and Dimmed, both known to activate fibroin gene expression in silkworm^67,68,71,72^ ; Vsx1, a homeobox TF that binds the *FibH* promoter directly in silkworms^47^ ; as well as Midline and Sine oculis, for which a function in the PSG remains to be studied. Among MSG-enriched TFs factors, Antp and Gsb are known to impact MSG identity and morphology in silkworms^28,29,44,73^. Previous research in silkworms has shown that Gsb is a key enforcer of ASG and MSG identity as it is responsible for both transcriptional repression of fibroin gene expression and symmetrical cell growth, necessary qualities for sufficient silk production and spinning^57,73,74^. We also detected additional genes encoding the TFs Bx-like, Vvl, Eyegone (Eyg), Poxn, Gsb-N, Dichaete (D), and Sox21b, that may contribute to the specialization of the MSG transcriptome.

Second, specialization of either region may involve enzymes that modify the secreted content. Pyralid sericins are water soluble, glue-like proteins that likely undergo post-translational modifications^12,75^. The gene *Pgant5* showed a strong enrichment in the MSG (**Fig. 3A-B**). In *Drosophila*, Pgant5 performs the first enzymatic step leading to the addition of N-acetylgalactosamine (GalNAc) sugar to Serine/Threonine (Ser/Thr) residues in the gut and salivary glands, a process known as mucin-type O-glycosylation^76,77^. While the *Plodia* major sericins show numerous Serine residues (**Fig. 3C-D**), and that sericins undergo GalNAc O-glycosylation in *Bombyx*^78^, Pgant5 is unlikely to modify sericins here as it lacks an N-terminal Signal Peptide (**Table S8**). Instead Pgant5 may thus glycosylate intracellular factors, akin to its role in fly intestinal cells^79^.

In contrast, the MSG enzymatic factors mentioned below each carry an N-terminal signal peptide (**Table S8**), indicating they are all targeted to secretory organelles and supporting a role in the modification of silk factors. The *ERp60* gene, encoding a disulfide isomerase enzyme, is highly expressed in the MSG and may be required for the formation of covalent bonds between silk factors such as FibL and FibH^22^. Similarly, we found MSG-enrichment of enzymes involved in the degradation of complex lipids such as several lipase genes (*Lip1*, *Lip3*, *Lip3-like*, *Lip4*, *LipF*, *LipK*, *LipK-like*), a glucocerebrosidase gene (*Gba1c*), and a saposin gene (*Sap-r*). This detection of lipid modifiers mirrors recent findings in *Galleria* moths and *Limnephilus* caddisflies^75,80^, suggesting this process is a key aspect of silk biology in Amphiesmenoptera (*i.e.* Lepidoptera and Trichoptera).

### Spatial HCR identifies two specialized spatial domains within the MSG

The RNAseq dataset depicts an overview of transcriptome divergence between two exocrine tissues, secreting either the silk fiber or its coating layers. The PSG transcriptome was consistent with a specialized secretory role focused on producing the fibroin core of the silk fiber. Meanwhile, the apparent transcriptomic complexity of the MSG could reflect either the presence of multiple distinct cell types, or greater molecular heterogeneity within a more homogeneous population. To assess this, we selected 11 MSG-enriched genes and profiled their spatial expression pattern using HCR.

Expression of the tandem-duplicate genes, *Ser1a* and *Ser1b*, were found to be restricted to the posterior region of the MSG directly adjacent to the anterior PSG as defined by *FibH* expression **(Fig. 4A-B)**. HCR also found expressions of *SerP150* and *Ser3a* are limited to the anterior region of the MSG **(Fig. 4A-B)**. In contrast, *Mucin12*, *MG4*, *Ser3b* (a *Ser3a* paralog), and *Zonadhesin1* were found to have expression throughout the entire MSG compartment **(Fig. 4C-D)**. These results suggest that the MSG consists of two cell types each restricted to two spatial domains: MSG-P, a small posterior compartment specialized in the expression of inner coating layer proteins (*e.g.* Ser1a/Ser1b) ; and MSG-A, a longer, more anterior compartment of the MSG that secretes major sericins (*Ser3a*/*Ser3b*, *MG4*, *SerP150*, *Mucin12*) and *Zon1*.

**Figure 4.**
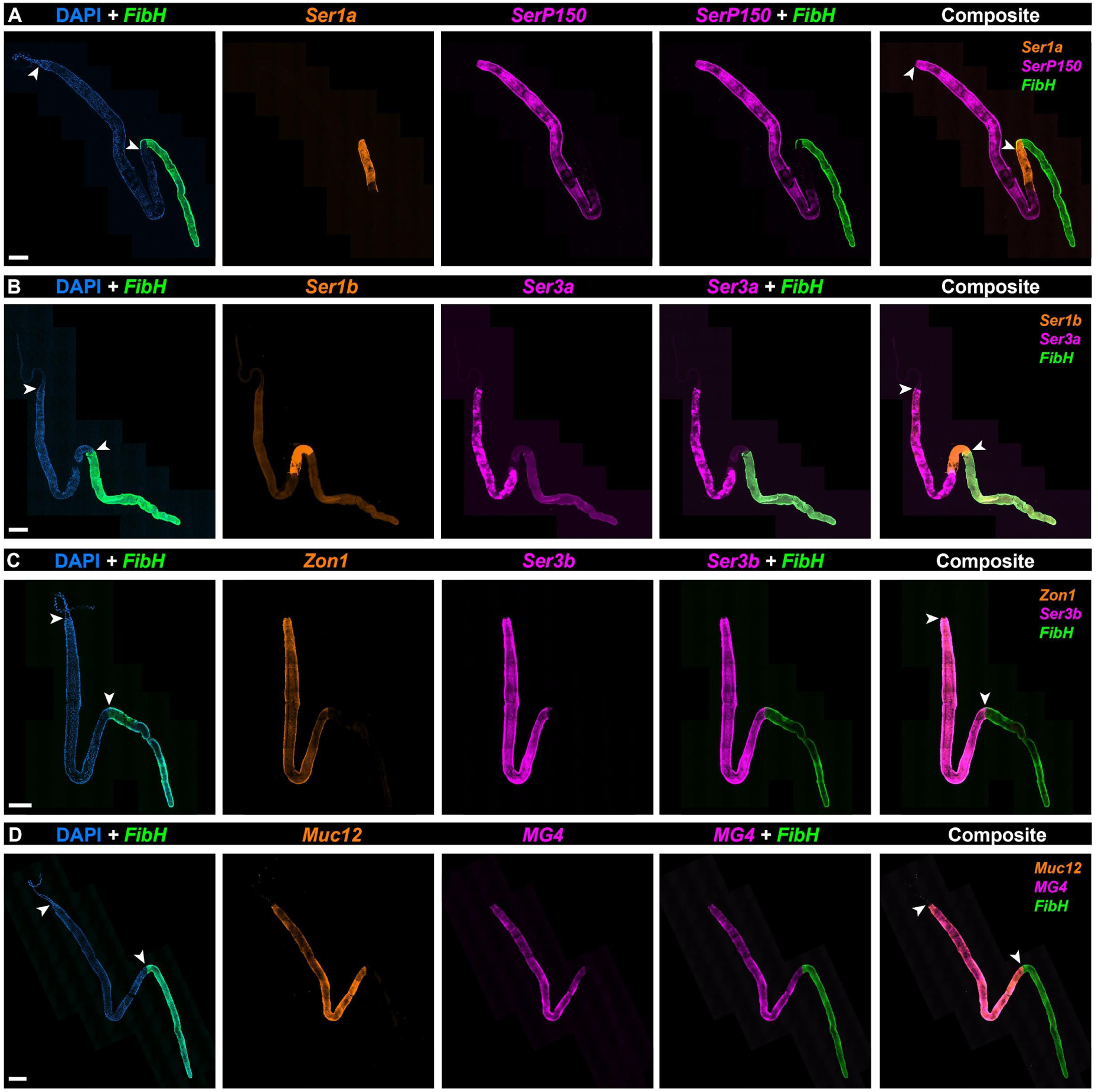
The MSG is compartmentalized into two subdomains of sericin and zonadhesin expression. Whole-mount silk gland HCR stainings of 8 MSG-enriched adhesive protein factors including 7 sericin factors (variably annotated as sericin and mucin genes) and a zonadhesin **A.** *Sericin1a* (*Ser1a*), *SericinP150* (*SerP150*), and *Fibroin Heavy chain* (*FibH*) mRNA. **B.** *Sericin1b* (*Ser1b*), *Sericin3a* (*Ser3a*), and *FibH* mRNA **C.** *Zonadhesin1* (Zon1), *Sericin3a* (*Ser3a*), and *FibH* mRNA **D.** *Mucin12* (*Muc12*), *MG4*, and *FibH* mRNA. Arrowheads: ASG/MSG (A-D: top) and MSG/PSG boundaries (A-D: bottom). Scale bars : A-D = 500 μm

### Expression of antimicrobial and conditioning factors

We broadly define silk conditioning factors as the molecules that are secreted by the silk gland to modify the properties of silk without playing a structural role. Of particular interest, seroins and serine protease inhibitors (serpins) act as broad-spectrum antibacterial and antifungal agents involved in preventing silk degradation^10,34,81^. In addition to the high expression of the seroin *Sn1*, RNAseq also detected the seroin gene *Seroin3* (*Sn3*) and the serpin *Silk protease inhibitor* (*Spi*) genes (**Fig. 3B-C**), previously studied in *Galleria* and *Ephestia*^10,75^. HCR assays show that *Seroin1* (*Sn1*) is expressed throughout the PSG and MSG, *Sn3* is high in the ASG and low in the MSG **(Fig. 5)**. While the function of these factors in microbial inhibition remains to be tested, gene expression patterns support the finding that *Plodia* silk contains bioactive antimicrobial properties^82^.

**Figure 5.**
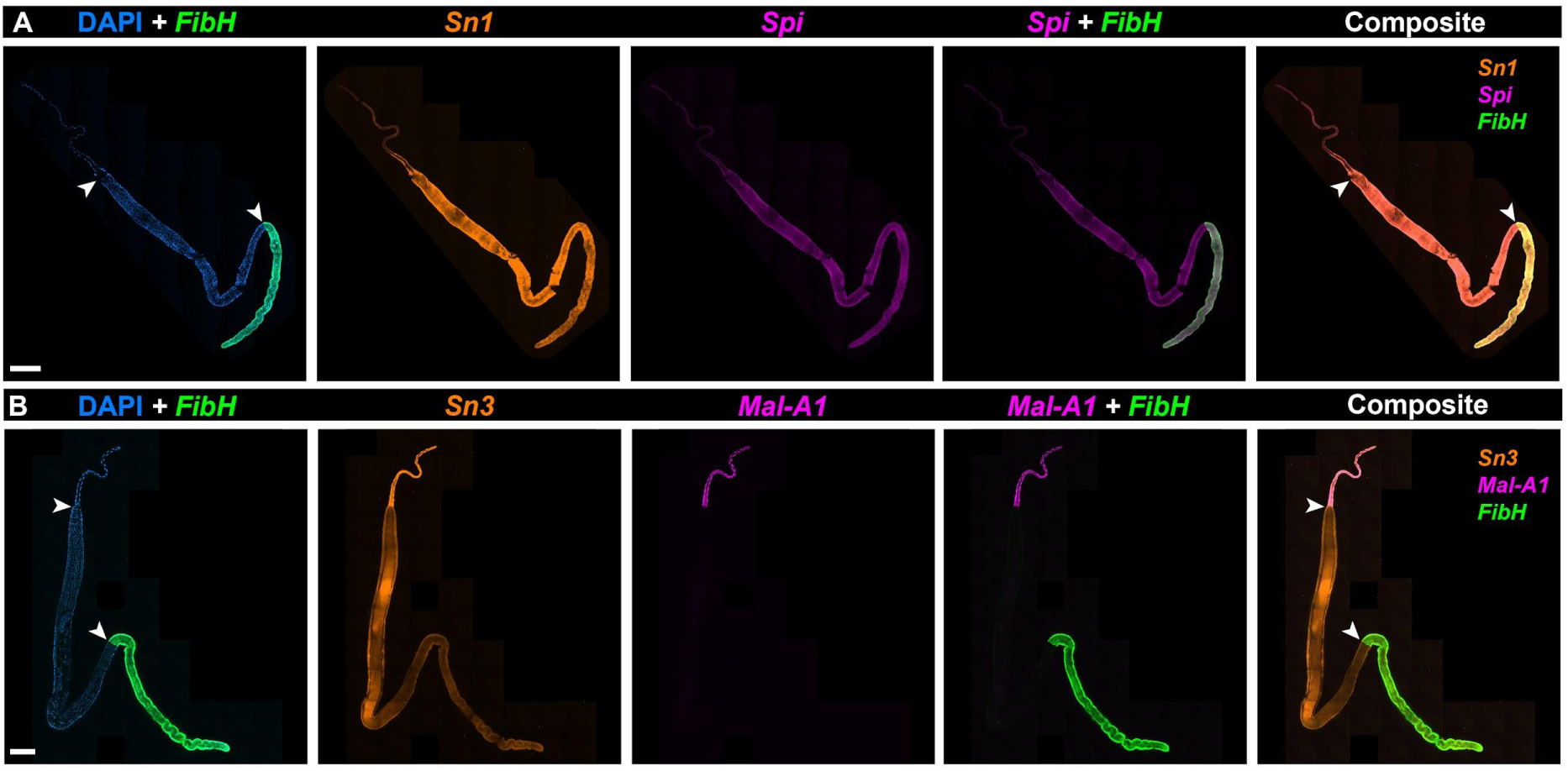
HCR detection of *seroins* and *Mal-A1* expression. **A.** Whole-mount silk gland HCR staining of *Seroin1* (*Sn1*), *Silk protease inhibitor* (*Spi*), and *FibH* mRNA. **B.** HCR staining of *MaltaseA1* (*Mal-A1*), *Seroin3* (*Sn3*), and *FibH* shown in a whole silk gland. Arrowheads: ASG/MSG (A-B: top) and MSG/PSG boundaries (A-B: bottom). Scale bars: A-B = 500 μm.

Lastly, RNAseq detected the expression of *Maltase-A1* (*Mal-A1*), a gene encoding an enzyme likely involved in the hydrolysis of maltose into glucose (**Fig. 3A-B**). As this putative function is intriguing, we sought to profile its expression with HCR, and found it to be specifically expressed in the ASG **(Fig. 5B**-**5C)**. This glucosidase enzyme includes an N-terminal signal peptide, suggesting it is actively secreted in the ASG lumen. We speculate that this enzyme may participate in glycoprotein maturation by modifying carbohydrates associated with silk proteins. Alternatively, it could be secreted into the outer silk coating, acting as a pre-digestive enzyme facilitating larval feeding in starch-rich environments.

## Discussion

### A roadmap for the gene expression analyses of insect exocrine tissues

As crucibles of biochemical innovation, insect exocrine tissues may offer an untapped reservoir of bioactive compounds of translational importance in biotechnology, medicine or agriculture. In this work, we used bulk RNAseq to profile gene expression in silk glands that were split at their approximated PSG/MSG boundary, and then refined the expression patterns of putative silk factors with a cellular spatial resolution, thereby identifying the major bioadhesive proteins secreted in the *Plodia* silk outer layer, as well as seroins that may confer its antimicrobial properties^34,35,82^. In combination with the availability of reliable transcriptome or genome annotations in a given species, this strategy offers a powerful way to decipher the molecular biology of exocrine glands across insects. Indeed, secreted genes can be identified by the presence of N-terminal signal peptides, and our data suggest that peptidic components of biological importance (e.g. fibroins and sericins) are expressed at high levels in secretory cells (**Fig. 6**). In Lepidoptera alone, more than 34 exocrine tissues have been described—most of them awaiting molecular characterization— including venom glands, myrmecophilic organs, and salivary glands that provide specialized interactions with predators, symbionts, and host plants^83^. In addition, other holometabolous insects such as wasps, sawflies, and glowworms also use labial glands to produce diverse silks that differ in composition with the lepidopteran silk^1^; gene expression profiling combining RNAseq and HCR could depict the modalities of convergence in the molecular toolkits underlying silk production across independently evolved insect lineages.

**Figure 6.**
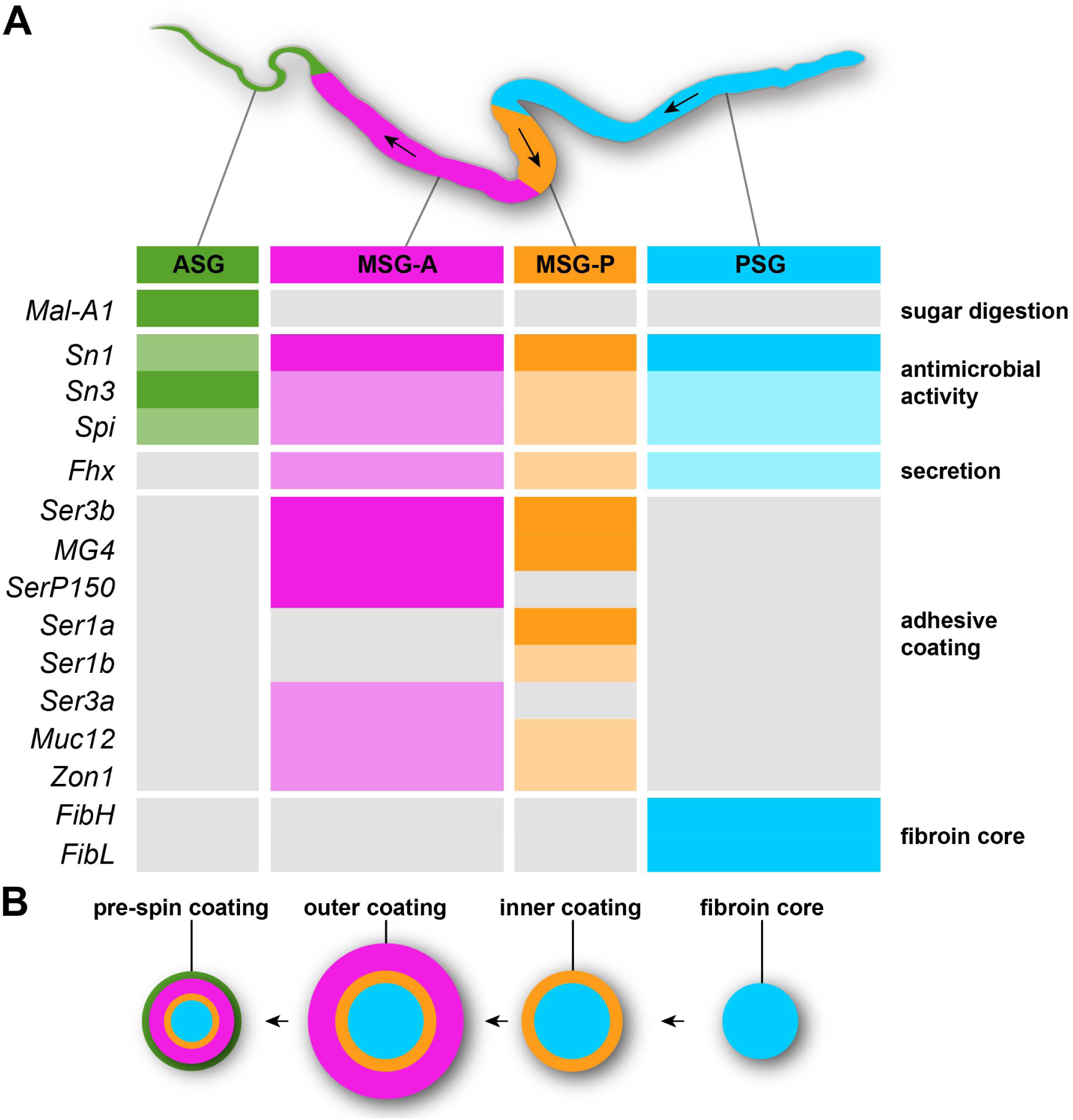
The regionalized expression of major *Plodia* silk factors define four specialized exocrine tissues. **A.** Summary of the HCR expression data for 11 silk factors. Color rectangles highlight regions of expression (full color : strong HCR signal, or TPM > 10,000 in MSG/PSG; dimmed colors : faint HCR signal or TPM < 10,000) ; grey rectangles indicate low expression (no HCR signal, or TPM < 2.5 in MSG/PSG). Arrows : direction of silk processing. **B.** Schematic view of the silk fiber, with successive layers of coating (from right to left) secreted along the specialized silk gland regions.

### Secretory specializations and cell-type regionalization of the silk gland

We generated deep transcriptomes of the *Plodia* last instar MSG and PSG and uncovered the identity and spatial domains of major silk factors, including fibroins, sericins, and seroins (**Fig. 6A**), some of which are sequentially secreted to constitute the concentric layers of the silk fiber and its coating (**Fig. 6B**). These data suggest the *Plodia* silk gland mainly consists of four regions and exocrine cell types: PSG cells specialized in core fibroin secretion, two successive types of MSG cells each secreting a distinct mixture of adhesive proteins, and ASG cells.

In addition, we started to uncover the regionalized expression of antimicrobial factors such as seroins and serine-protease inhibitors, suggesting that the deposition of silk conditioning factors can be assigned to distinct layers of the fiber. In conjunction with previous expression studies in other lepidopteran silk glands^10,34,36,75,84,85^, these data suggest that the expression of bioactive molecules involved in the prevention of silk degradation is an important conserved quality of lepidopteran silk. Future studies of this process could extend to other seroins and serpins detected in the dataset, and assess the effects of larval stage and microbial exposure to expression patterns.

Unexpectedly, HCR spatial assays identified *Mal-A1* as a specific marker of the ASG. This result highlights a limitation of our RNA-seq study: because silk glands were sectioned at the MSG/PSG boundary, these samples also included the comparatively smaller ASG gland, and some of the MSG-enriched transcripts described in this study may turn out to be enriched in the ASG instead. Due to the relative small size of the ASG tissue compared to the rest of the MSG (**Fig. 1**), we can infer that ASG-specific transcripts are largely obscured by the MSG contribution to this dataset, and further studies of the elusive functions of the ASG would benefit from RNAseq resampling combined with HCR spatial assays.

### Spatial homology between silk gland sub-domains across Lepidoptera

The overall morphology and exocrine function of silk glands appears conserved across silk-spinning lepidopterans, with a PSG portion dedicated to the secretion of the fibroins FibL and FibH, and an MSG portion involved in the coating of outer layers of adhesive proteins. RNAseq and HCR spatial analyses of silk factor expression confirm this big picture in *Plodia* (**Fig. 6**), complementing previous analyses in other pyralid moths^10,38^. We found that the tandem duplicate genes *Ser1a* and *Ser1b* are specifically expressed in a short domain, previously known as the rear-MSG in Pyralidae^10^. Using qPCR in silk gland sections of the pyralid *Ephestia*, Wu *et al.* previously found a similar pattern with both *Ser1a* and *Ser1b* showing rear-MSG enrichment, and proposed that *Sericin1* genes, characterized by a signature CxCx motif in their C-terminal region, highlight the conservation of this compartment with *Bombyx* and beyond^28^. Microsynteny analyses further support the homology of *Plodia Ser1a* and *Ser1b* with *Bombyx Ser1*, and the HCR mRNA expression of *Plodia Ser1a* and *Ser1b* was also reminiscent of the spatial expression of *Ser1* in *Bombyx* fourth instar larvae, in a subsection of the MSG immediately anterior to the PSG that later extends to the whole MSG during the fifth instar^28,29^.

In *Plodia*, restricted expressions of *SerP150* and *Ser3a* define a long, anterior domain of the MSG (here dubbed MSG-A), while both MSG-A and MSG-P expressed *Ser3b*, *MG4*, *Muc12*, and *Zon1*. We did not find molecular evidence that this MSG-A region can be subdivided into two portions, as proposed for *Galleria* wax moths^75^, and as this is the first time that these adhesion genes are characterized in whole-mount expression assays, further investigation is needed to assess whether this sub-regionalization applies to other species.

In summary, we can affirm that the lepidopteran silk glands share a homologous PSG, a *Ser1*-positive MSG-P compartment involved in the secretion of the silk inner coating layer ^10,28,29^, and an MSG-A region involved in the secretion of the outer coating layer, marked by spatially restricted domains of *SerP150* and *Ser3a* in *Plodia*. Of note, sericin expression and alternative splicing profiles can vary across stages in the silkworm, as cocoon-specific silk differs from the silk spun during earlier larval stages^86,87^. In the future, HCR mRNA *in situ* hybridizations should be a valuable tool to assess evolutionary variations across Lepidoptera, including stage-specific differences in sericin usage.

## Acknowledgements

We thank Donya Shodja and Christa Heryanto for their support rearing *Plodia*; Luca Livraghi for guidance on HCR protocol optimization; Patricia Hernandez and Aleksandar Jeremic for providing access to confocal microscopes; Anastas Popratiloff and personnel of the GW Nanofabrication and Imaging Center (GWNIC) for support with spinning disk confocal microscopy. Micro-computed tomography was conducted at the Research Service Centers of the Herbert Wertheim College of Engineering at the University of Florida with the valuable assistance of Gary Scheiffele. This work was funded through a collaborative IntBIO grant from the National Science Foundation, (awards MCB-2217156 to AM and MCB-2217159 to WLS), and a National Institutes of Health National Institute of General Medical Sciences Maximizing Investigators’ Research Award to WLS (NIH NIGMS R35-GM147041). Microscopy equipment at the GWNIC was funded by the award NIH 1S10OD010710-01.

## Data Availability

Detailed procedures for immunofluorescence, fluorescent dye, and HCR stainings can be found on the Open Science Framework repository^88^. The raw reads from RNA sequencing have been deposited in the NCBI Sequence Read Archive under BioProject PRJNA1241317. All code associated with this project can be found in the GitHub repository: https://github.com/jasalq/Plodia_Silk_RNAseq.

## Methods

## KEY RESOURCES TABLE

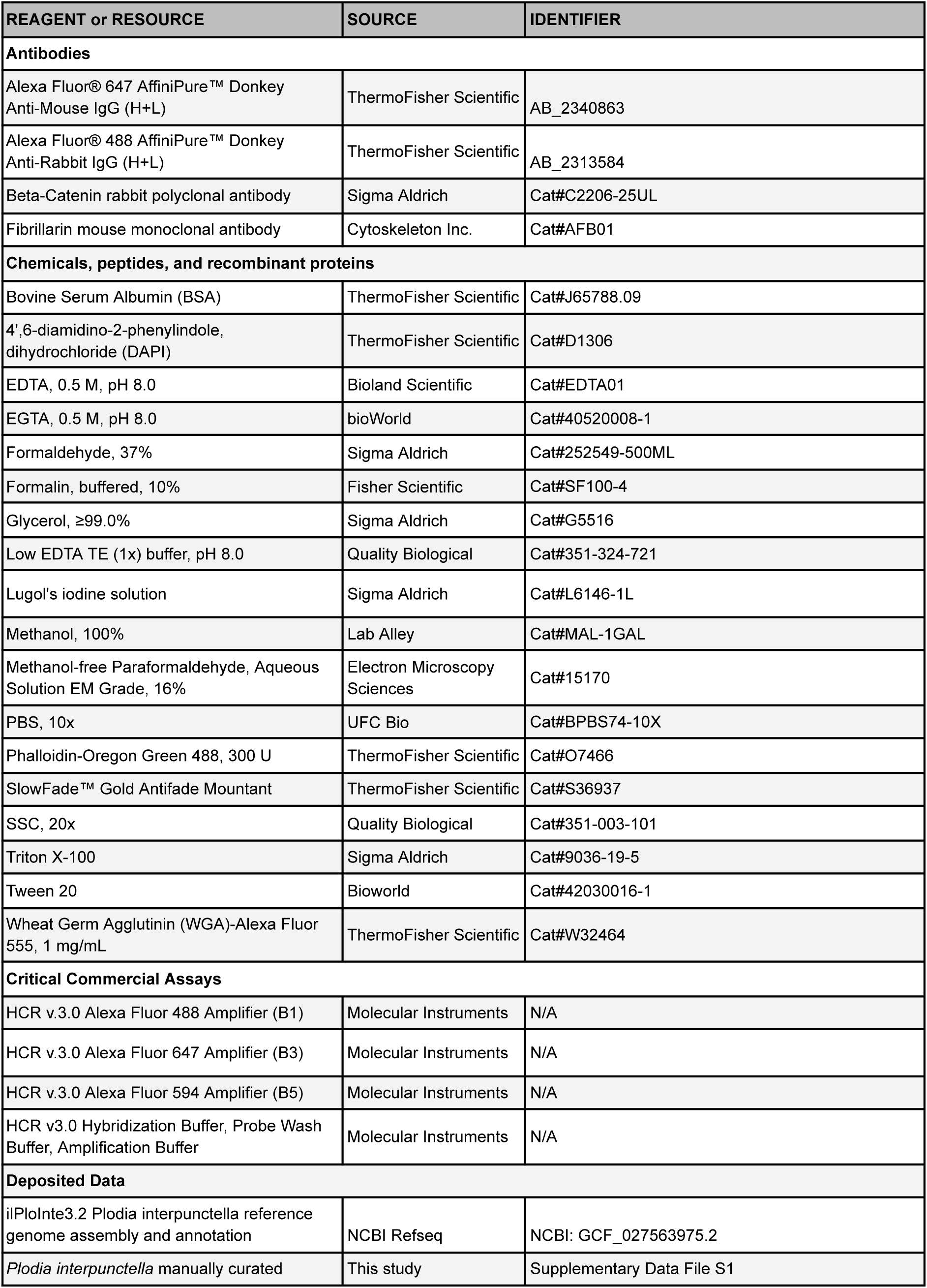
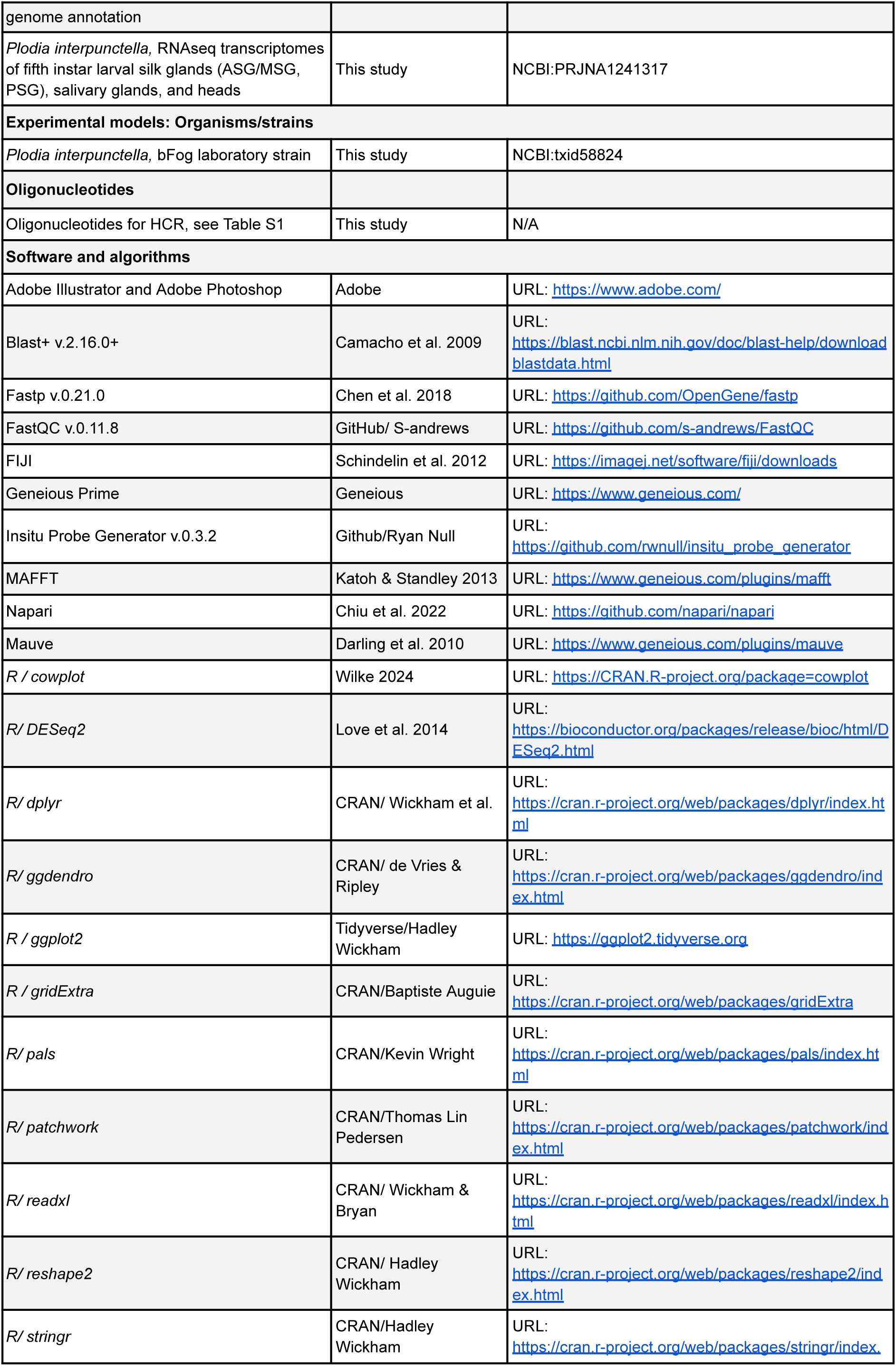
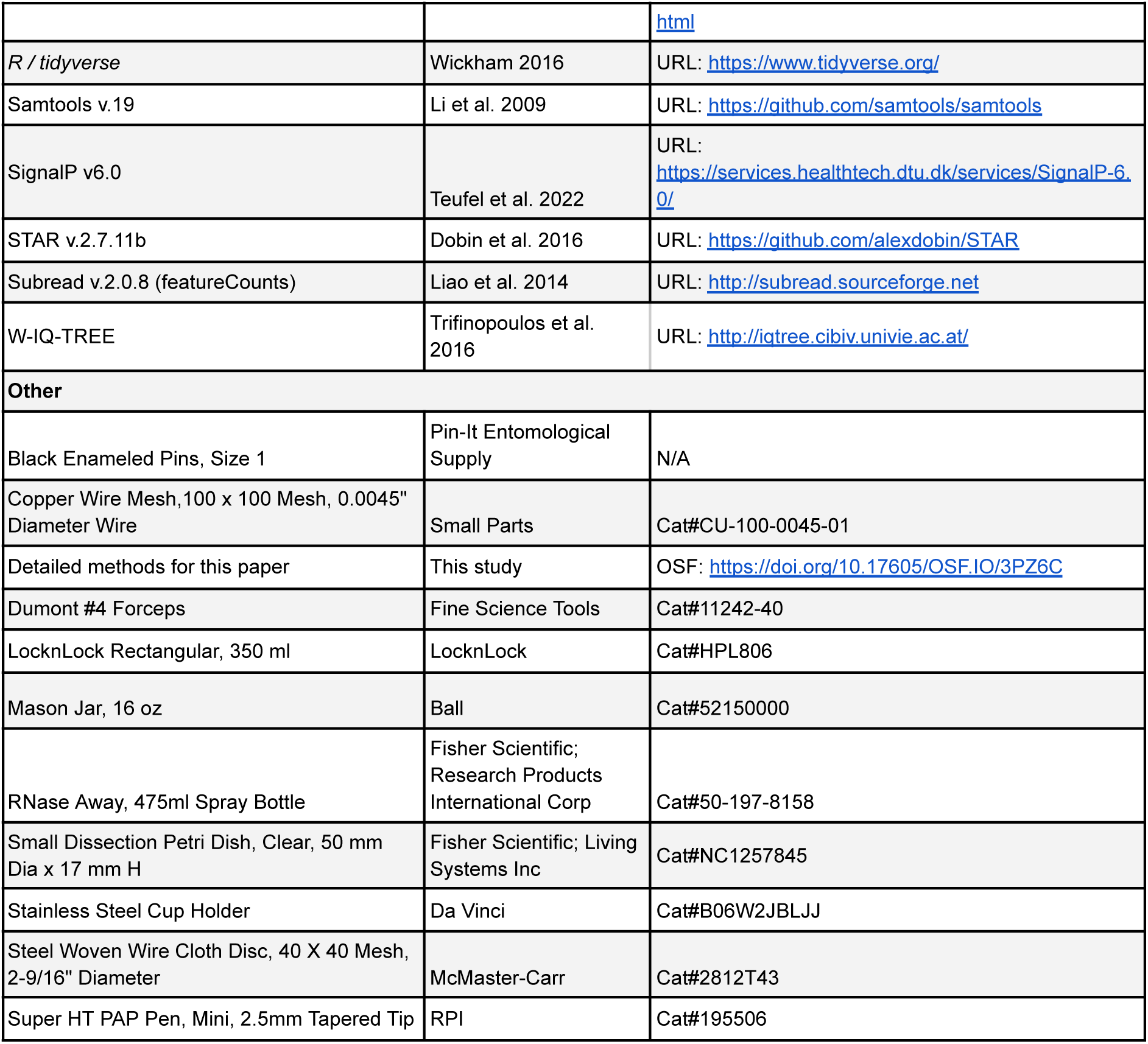

## EXPERIMENTAL MODEL AND STUDY PARTICIPANT DETAILS

The wild-type *bFog* laboratory strain of *P. interpunctella* was reared in the laboratory from egg to adulthood in a growth chamber at 28°C with 60-80% relative humidity and a 14:10 h light:dark cycle^53^. Rearing methods are available online on the Open Science Framework (OSF) repository^88^. Briefly, egg laying was induced by CO_2_ narcosis of adult stock in an oviposition jar, and a weight boat containing 10-12 mg eggs was placed in a rearing container containing 45-50 g of wheat bran diet with 30% glycerol. At 28°C, this life cycle spans 29 days from fertilization to a reproductively mature adult stock.

## METHOD DETAILS

### Micro-Computed Tomography

A fifth instar larva was fixed in 10% phosphate buffered formalin overnight before the skin layer was nicked by needle pins and then returned to fixative for 48 hours. Fixed larvae were submerged in Lugol’s iodine contrast solution for 1 week and rinsed in ultrapure water to remove excess contrast solution directly prior to scanning. The stained specimen was scanned on a GE Phoenix v|tome|x m 240 CT scanner (GE HealthCare Technologies, USA) equipped with a 180 kV transmission tube and diamond target with a 4.98 μm voxel size, 120 μA current, and 80 kV voltage. 3D volume files were analyzed and rendered with Volume Graphics VGStudio Max v2024.2 software suite (Volume Graphics, GER). Silk gland segmentation, refinement, and smoothing was completed with VGStudio Max’s segmentation tools on reconstructed 3D volumes.

### Antibody and fluorescent dye stainings

Protocols for the dissection, immunofluorescent staining, and dye stainings of *Plodia* silk glands are described online on the OSF repository^88^. Briefly, whole silk glands were dissected from wandering fifth instar larvae. For both immunofluorescent and dye stainings the samples were fixed in a 4% methanol free formaldehyde solution for 20 min and then washed three times for 5 min in PBT at room temperature. For immunofluorescent staining, the samples were washed for four 5 min washes in PT and then blocked in a PT-BSA solution at room temperature. Samples were incubated in the primary antibodies solution overnight at 4°C. After incubation two quick washes and three 8 min washes in PT were performed. Samples were then blocked again in PT-BSA for 10 min at room temperature before the secondary antibody solution was added for incubation at room temperature for 2 hours. Final washes in PT (two quick and four 15 min washes) were performed before a solution of 50% glycerol with 1.0 μg/mL DAPI (pH 7.4) was added to incubate either for 30-60 min at room temperature or overnight at 4°C. Samples were mounted in 70% Glycerol (diluted in PBS with pH 7.4) and imaged.

For dye stainings, after the samples were originally washed in PBT, a diluted solution of WGA-Alexa 555 in 1X PBS was added to the samples and left to incubate for 1 to 2 hours in the dark at room temperature. Three 5 min washes in PBT and two quick washes with PT were then performed before staining in a diluted solution of Phalloidin (Oregon Green)-Alexa 488 in PT for either 2 to 4 hours at room temperature or 18 to 24 hours at 4°C. After staining, three quick washes and three 10 min washes with PT were performed at room temperature and a solution of 50% glycerol with 1.0 μg/mL DAPI (pH 7.4) was added to incubate either for 30-60 min at room temperature or overnight at 4°C. Samples were mounted in 70% Glycerol diluted in PBS under a #1.5 glass coverslip before imaging.

### Fluorescent In situ Hybridization Chain Reaction (HCR)

A modified version of the third-generation *in situ* hybridization chain reaction protocol was performed^65,89^, and is fully described with solution recipes on the OSF repository^88^. Briefly, probes were designed for target genes using the *insitu_probe_generator* software^90^. Silk glands were dissected in cold 1X PBS and fixed in a 3.7% formaldehyde solution for 20 min at room temperature. After four 5 min washes in PBT the samples were dehydrated progressively in cold methanol diluted in PBS until a final concentration of 100% methanol was reached. Some samples were stored in 100% methanol at -20°C before continuing the protocol. All samples, freshly dehydrated or stored, were rehydrated progressively by adding decreasing concentrations of cold methanol diluted in PBS. After three 5 min washes in PBT, samples were placed in 200 μL of pre-warmed probe hybridization buffer and placed at 37°C for 30 min while shaking. Samples were then incubated while shaking overnight at 37°C while in probe hybridization solution. After the incubation, four 15 min washes in probe wash buffer were performed at 37°C. At room temperature, three 5 min washes in 5 x SSCT were performed while shaking and samples were pre-amplified in amplification buffer for 30 min-1 hr. During incubation, the amplifier solution was prepared by heating 4 μL of each amplifier for 90 sec at 95°C and allowing them to cool for 30 min at room temperature in the dark. The amplifiers were then added to 200 μL of amplification buffer and after the old amplification buffer was removed, the amplifier solution was added to the samples. Samples were incubated overnight in the dark at room temperature while shaking. Amplifier solutions were then removed from the samples and stored at -20°C for re-use. Samples were washed with 200 μL 5 x SSCT at room temperature for two 5 min washes, two 30 min washes, and one 5 min wash. After removal of solution from the last wash, samples were incubated in a solution of 50% glycerol with 1.0 μg/mL DAPI (pH 7.4) either for 30-60 min shaking at room temperature or overnight shaking at 4°C. After removal of the DAPI staining solution, samples were mounted in 70% Glycerol diluted in PBS or SlowFade Gold Antifade Mountant and imaged.

### Confocal microscopy and image processing

Whole-gland fluorescent microscopy images were obtained with a Zeiss Cell Observer Spinning Disk confocal microscope mounted with a 10x objective (Plan-Apochromat, 0.45 NA), allowing the rapid acquisition of stitched images across the whole tissue. Fluorescent images acquired from the Spinning Disk confocal were pre-processed using the Zen acquisition software using a shading reference approach to correct for tiling artifacts. High-magnification fluorescent microscopy views of silk glands were obtained. Stacked acquisitions were also obtained on an Olympus FV1200 confocal microscope and a Zeiss LSM 800 confocal microscope, each mounted with mounted 20x and 60x objectives. Fluorescent acquisitions were processed in FIJI and Napari^91,92^. Adjustments of contrast limits were applied independently to each fluorescent channel. Brightfield images were acquired using a Nikon D5300 camera mounted to a Nikon SMZ800N trinocular dissecting microscope, equipped with a P-Plan Apo 1X/WF 0.105 NA 70 mm objective.

### RNA Sequencing

Tissues were obtained from fifth instar wandering larvae of the *Plodia bFog* strain^53^. Silk glands, salivary glands, and larval heads were dissected in cold 1X PBS. Silk glands were sectioned into two segments by cutting at the boundary of the MSG-P and PSG, as defined by expression of *FibH* and *FibL* (**Fig. 1A**; **Fig. 1E-F**). Each tissue type was obtained from two individuals and then pooled into 2 mL tubes containing 500 μL of TRI Reagent (Zymo Research) which were stored at -80°C. This process was repeated so that there were four biological replicates for each of the four tissue types. Samples were sent for total RNA extraction, poly-A enriched library preparation, and PE150 sequencing on an Illumina NovaSeq X instrument with a target yield of 30M reads per library, outsourced to Genewiz (South Plainfield, NJ), PSG samples were sequenced twice due to a suboptimal yield in the first sequencing run, and the resulting technical replicates were combined using the *collapseReplicates* command in DESeq2. Quality of the RNA sequencing data was accessed using FastQC v.0.11.8^93^. Adapters and PolyG tails were trimmed using *Fastp* v.0.21.0^94^ with the options *–-detect_adapter_for_pe* and *–trim_poly_g*. *FastQC* was run post-trimming to check for adapter and polyG contamination and assess post-trimming read quality. RNA sequencing reads were aligned to the *P. interpunctella ilPloInte3.2* reference genome (GCF_027563975.2) using *STAR* v.2.7.11b^95^. STAR alignment was repeated with the IntronMotif output from the original run to better resolve splice junctions prior to read counting.

### Functional annotations of *Plodia* gene names

The *ilPloInte3.2* NCBI RefSeq annotation of the *P. interpunctella* genome (*GCF_027563975.2-RS_2024_04*) was used for RNAseq transcript mapping resource with minor modification: two consecutive transcripts (XM_053766583.1 and XM_053766584.1) were merged to form the retained *Pi_Ser1b* gene, and we added or corrected the gene models for a total of 23 antimicrobial peptide genes using manual validations of expression evidence, signal peptides, and sequence analysis. Of note, the *ilPloInte3.2* NCBI RefSeq annotation contains automated gene and protein names that are biased towards vertebrate gene nomenclature. We took two steps to complement this functional annotation with names that would be biologically informative in analyses of insect transcriptomes. First, we curated and edited the name of 30 silk factor genes, using a nomenclature reflecting direct homology to silk factors from other pyralids species as shown in this study (**Table S7; S9**) and previous publications^10,69^. Second, we also added functional annotations based on sequence similarity to *Drosophila melanogaster* genes to enrich gene names with biological knowledge from a reference insect model organism. To do this, the *ilPloInte3.2* RefSeq_protein sequences were extracted using NCBI Batch Entrez, and a BLASTp reciprocal homology search using the BLAST+ tool was performed with the translated protein sequences from the FB2025_02 Flybase release of the *D. melanogaster* genome annotation^96^. These imputed names can be seen in the “Symbol” column of **Table S9 and File S1**. Read counting was performed using *FeatureCounts* (Subread v.2.0.8)^97^ using the manually curated *ilPloInte3.2* NCBI RefSeq annotation, and gene feature list with manually edited names, respectively provided online in the GTF and TSV formats (**File S1; Table S9**).

### Sericin and seroin genes microsynteny analyses and sequence alignments

Sericin and seroin gene synteny analyses were initially conducted using the *progressiveMauve*^98^ plugin implemented in Geneious Prime (2023.0.2), using the corresponding NCBI RefSeq genomic scaffolds for *B. mori* (NC_085117), *G. mellonella* (NW_026442003), and *P. interpunctella* (NC_071322). Once the target regions were identified, homology of individual genes was tested between *P. interpunctella* and putative *B. mori* or *G. mellonella* syntenologs using reciprocal TBLASTN searches using predicted protein queries from one species, to both *RefSeq_rna* and *RefSeq_genomes* of the other species. The resulting microsynteny relationships were then manually curated in Adobe Illustrator visualizations of the annotated genomic intervals. Homologous genes of additional species were obtained from the literature^10,23,41,69^ or using NCBI TBLASTN against Lepidoptera RefSeq transcriptomes, and aligned with MAFFT ^99^ in Geneious Prime. Accession numbers for the homologous genes used can be found in **Table S7**. Additionally the seroin amino acid alignment matrix generated by MAFFT was used by IQ-TREE with default parameters to build a phylogenetic tree^100^.

## QUANTIFICATION AND STATISTICAL ANALYSIS

### Differential Gene Expression Analyses

The count data generated by FeatureCounts was used to perform differential expression analysis using DESeq2 in RStudio^101^. The DESeq2 function *collapseReplicates()* was used to combine sequencing runs for the PSG samples prior to performing the analysis. The experimental design *∼Tissue* was used to define the four tissues of interest (MSG, PSG, head, salivary gland), each with four biological replicates. The initial set of differential expressed genes (DEGs) in the MSG vs. PSG contrast is defined by an adjusted *p* < 0.05 (**Table S2**). In parallel, TPM (transcripts per million) values were calculated for each gene to measure relative transcript abundance within each biological replicate^70^. First, the read counts from the unormalized count matrix were normalized by gene length in kilobases, as calculated by FeatureCounts, to obtain an RPK (reads per kilobase) value for each gene. TPM values were then calculated per sample with the formula: *TPM* = *RPK/Scaling factor* × 1*e*6, where the scaling factor corresponds to the sum of all RPK values within the sample. Mean TPM values for each tissue were then obtained by averaging TPM values per gene across replicates.

The R package ggplot2^102^ was used to produce a scatter plot of gene expression divergence between tissues and relative abundance within tissues (log_2_FoldChange and log2TPM). The ggplot2 package was also used to produce a heatmap plot featuring the top 229 DEGs between MSG and PSG clustered by gene expression profile similarity with a significance threshold of an adjusted *p* < 0.01 and a | log_2_FoldChange | > 3, following a variance stabilizing transformation using the function *vst()* from DESeq2. After the variance stabilizing transformation the expression data was collapsed by tissue and scaled to a z-score. Expression profiles by gene were clustered using hierarchical clustering by the R package *ggdendro*^103^, and *ggplot2* was then used to visualize the final heatmap. N-terminal signal peptides among top PSG and MSG enriched genes were predicted using SignalP v6.0^104^.

## ADDITIONAL RESOURCES

Detailed fluorescent dye, immunofluorescence, HCR staining, and dissection protocols can be found on an Open Science Framework methods repository^88^. The manually curated *ilPloInte3.2* NCBI RefSeq annotation GTF file and gene list can be found as **File S1** and **Table S9**, respectively. Detailed tables of the results from the differential expression analyses can be found in **Tables S2-S6**. All code associated with this study can be found in a GitHub repository: https://github.com/jasalq/Plodia_Silk_RNAseq.

## Author Contributions

Conceptualization, Writing – original draft: J.D.A. and A.M. Investigation: J.D.A., M.B., L.E.E, W.L.S. and A.M. Formal analysis: J.D.A. Methodology: J.D.A and M.B. Visualization: J.D.A., M.B., L.E.E, and A.M. Funding acquisition and Supervision: W.L.S. and A.M. Writing – review & editing : L.E.E and W.L.S.

## Supplementary Information

**Figure S1.**
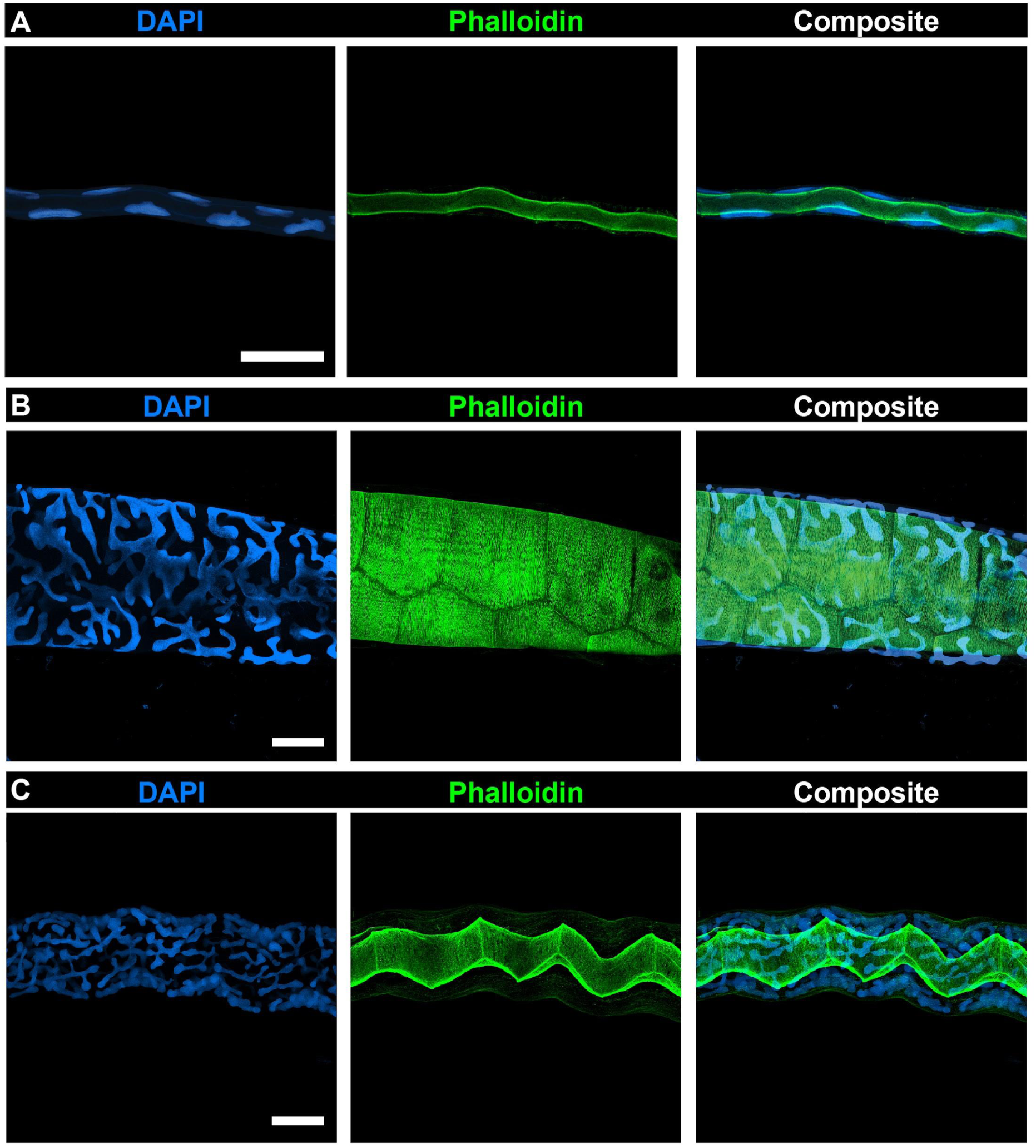
DAPI and Phalloidin staining reveal striking differences in cellular morphology and arrangement between compartments. **A-C.** DAPI and phalloidin staining of an ASG (A), MSG (B), and PSG (C). See <u>Video S3</u> for a video exploring the 3D projections of these images. The images from panel A-B are taken of the same gland while the images in panel C are taken of a different gland from an individual of the same developmental stage (wandering fifth instar). Scale bars: **A-C** = 100 μm

**Figure S2.**
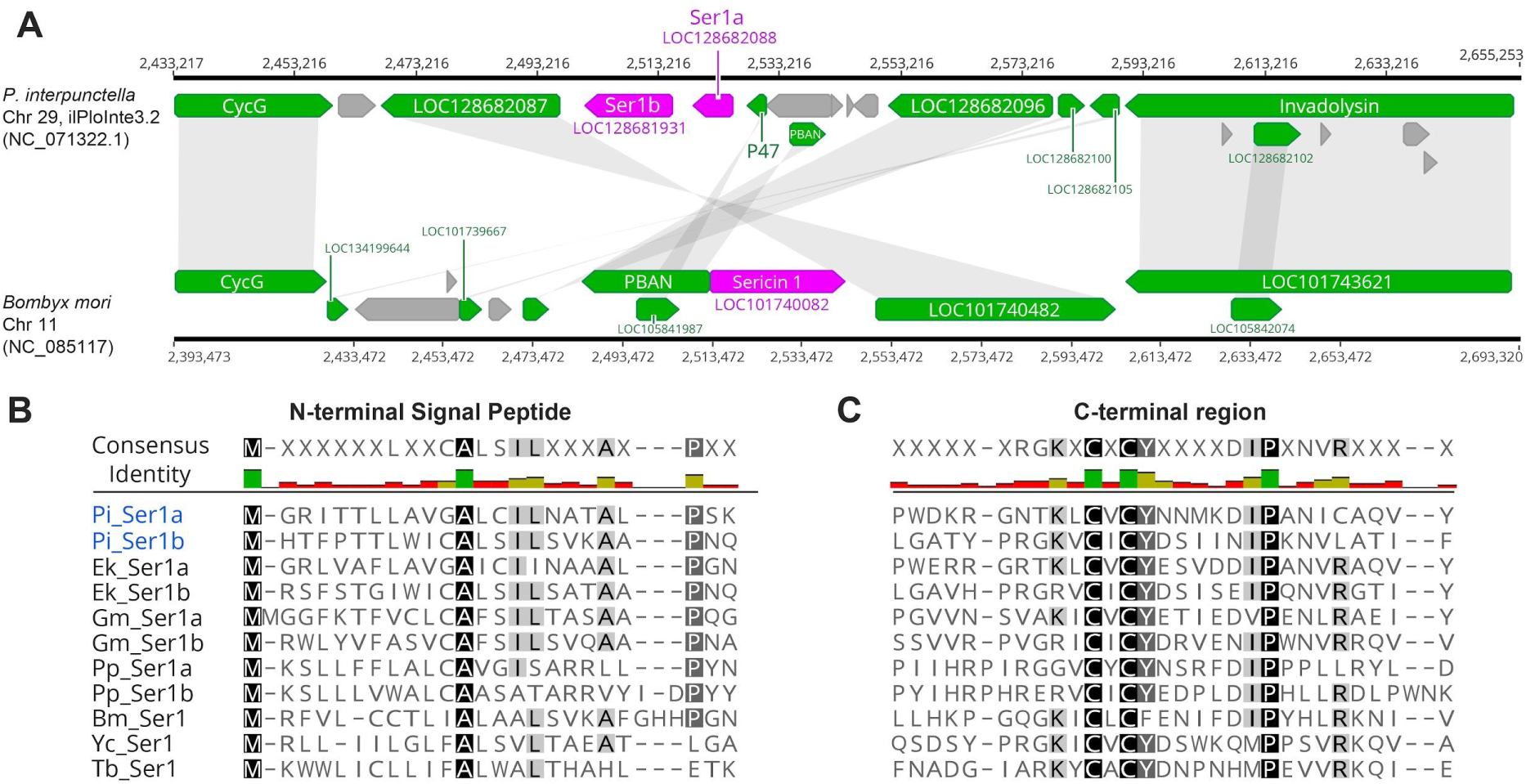
Homology of *Ser1* sericin gene copies across Lepidoptera. **A.** Synteny analysis reveals *Ser1a* and *Ser1b* genes of pyralid genomes (here on top, *P. interpunctella*) are found as tandem duplicates, within an inverted syntenic block containing the *B. mori Ser1* homolog (magenta). Grey fields indicate sequence matches using reciprocal TBLASTN between the predicted protein of a first species and the NCBI RefSeq_RNA dataset of the second species. **B-C.** Protein alignments of Ser1 syntenologs from genome annotations of lepidopteran species, largely based on previous analyses ^10,41,105,106^, with a focus on the N-terminal signal peptide (B) and the C-terminal region (C). The C-terminal CxCx motif is unique to this sericin orthology group. Gene and protein identifiers are listed in **Table S7**. Ek : *Ephestia kuehniella* (Pyralidae, Phycitinae), *Plodia interpunctella* (Pyralidae, Phycitinae); Gm : *Galleria mellonella* (Pyralidae, Galleriinae); Pp : *Pseudoips prasinana* (Nolidae; Chloephorinae); Bm : *Bombyx mori* (Bombycidae, Bombycinae); Yc : *Yponomeuta cagnagella* (Yponomeutidae; Yponomeutinae) ; Tb : *Tineola bisselliella* (Tineidae; Tineinae).

**Figure S3.**
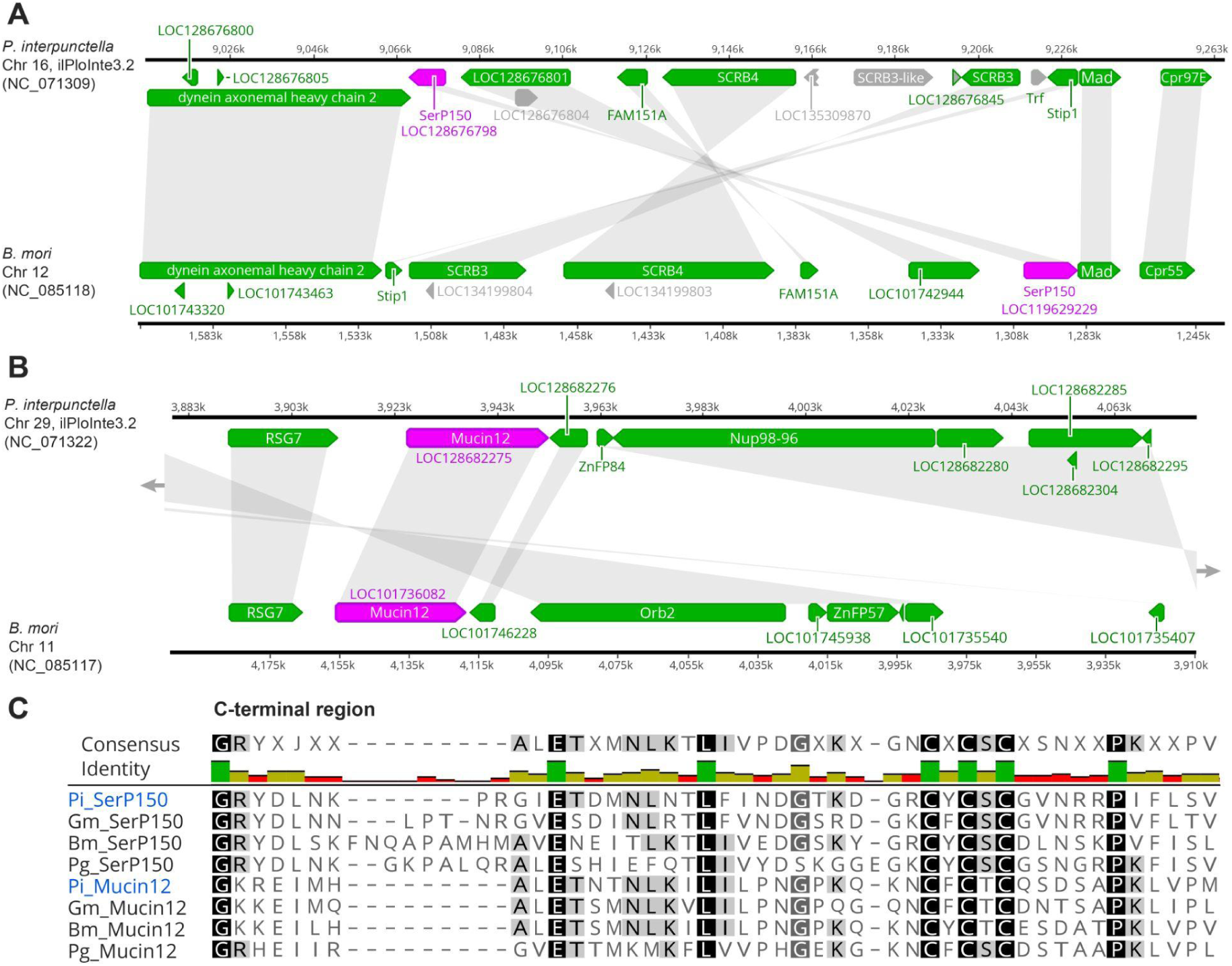
Homology and microsynteny of *SerP150* and *Mucin12* sericin gene orthologues in Lepidoptera. **A-B.** Microsyntenic relationships of the *SerP150* (A), based on a previous study^69^, and *Mucin12* (B) gene regions between *P. interpunctella* (top) and *B. mori*. Grey fields indicate sequence matches using reciprocal TBLASTN between the predicted protein of a first species and the NCBI RefSeq_RNA dataset of the second species. **C.** Protein alignments of SerP150 and Mucin12 syntenologs from various lepidopteran insects, characterized by a conserved CxCxC motif in their C-terminal domains. This alignment replicates the findings of a previous study^69^. Gene and protein identifiers are listed in **Table S7**. Pi : *Plodia interpunctella* (Pyralidae); Gm : *Galleria mellonella* (Pyralidae); Bm : *Bombyx mori* (Bombycidae); Pg : *Pectinophora gossypiella* (Gelechiidae).

**Figure S4.**
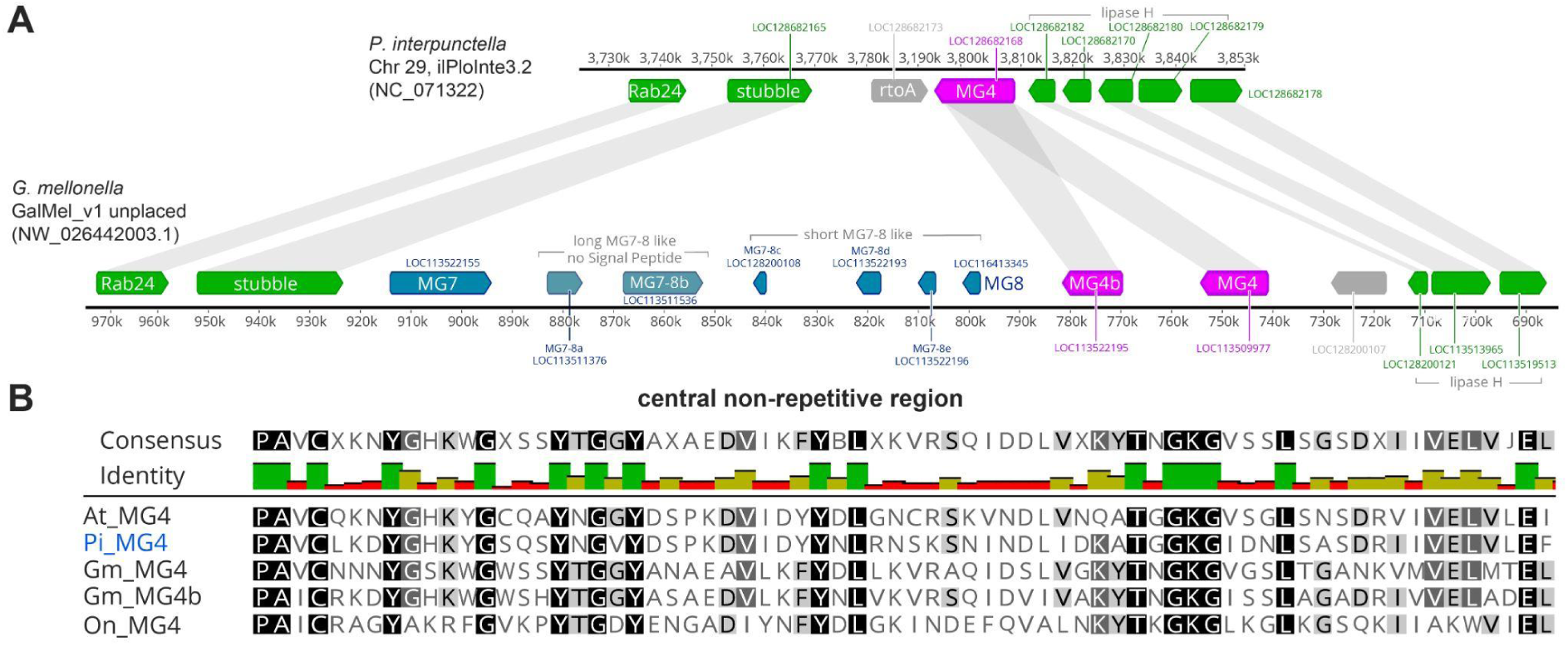
Homology of MG4 sericin factors in Pyraloidea. **A.** Synteny comparison between gene regions encompassing a single copy of the *MG4* sericin factor in *P. interpunctella* (top), and a cluster of related MSG-enriched genes specific to the *G. mellonella* genome, as previously described^10,12,107,108^. Grey fields indicate sequence matches using reciprocal TBLASTN between the predicted protein of a first species and the NCBI RefSeq_RNA dataset of the second species. *G. mellonella MG7-MG8* genes are likely paralog copies of *MG4*, consistent with an expansion of sericin genes in this species. **B.** Protein alignments of MG4 homologs from genome annotations of pyralid and crambid species (Pyraloidae), with a focus on the central domain, N-terminal to a serine-rich repeat region). Gene and protein identifiers are listed in **Table S7**. At : *Amyelois transitella* (Pyralidae, Phycitinae); Pi : *Plodia interpunctella* (Pyralidae, Phycitinae); Gm : *Galleria mellonella* (Pyralidae, Galleriinae); On : *Ostrinia nubilalis* (Crambidae, Pyraustinae).

**Figure S5.**
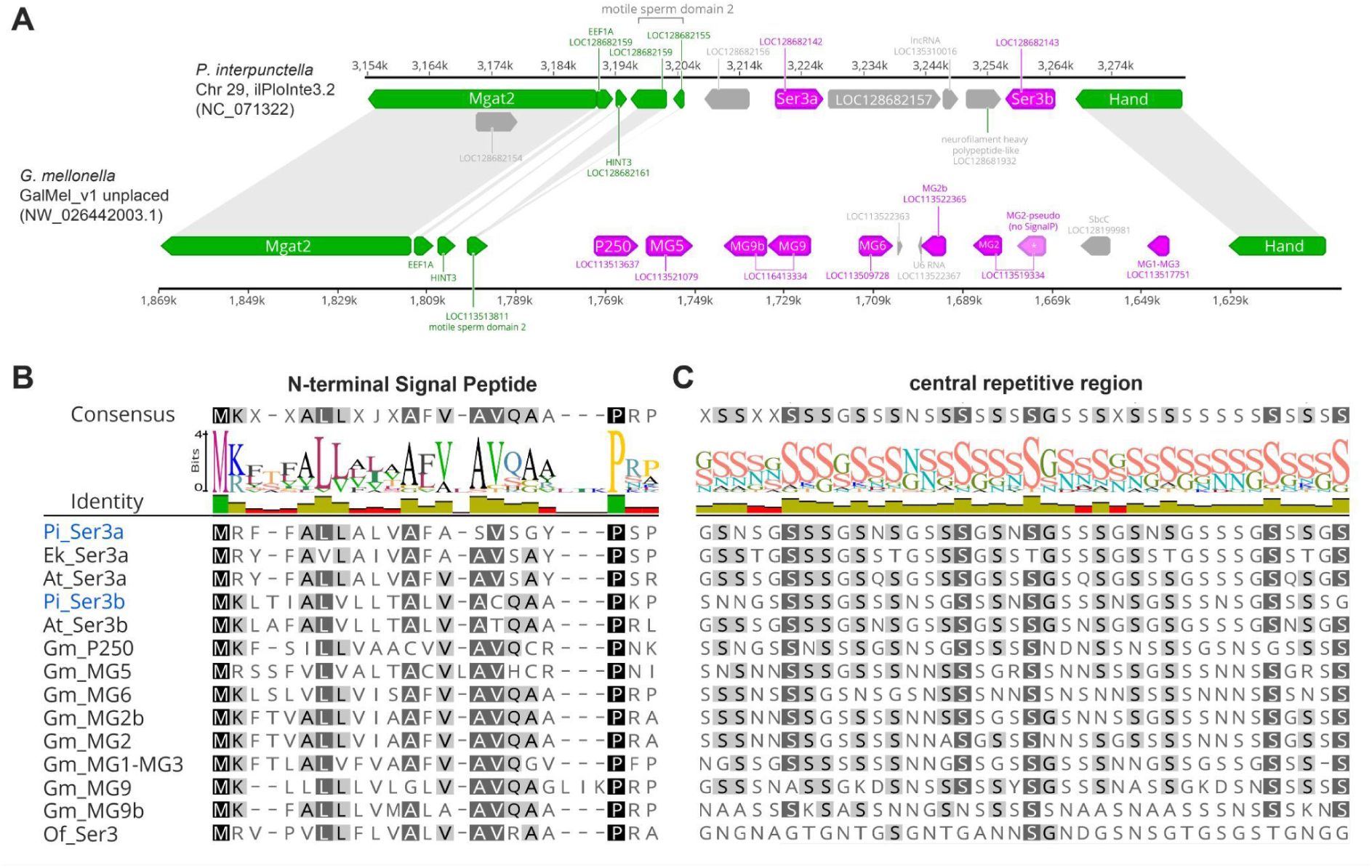
Homology of *Ser3* sericin factors in Pyraloidea. **A.** Synteny comparison between gene regions encompassing two copies of the *Sericin3* gene in *P. interpunctella* (top), hereby dubbed *Ser3a* and *Ser3b*, and a cluster of related MSG-enriched genes specific to the *G. mellonella* genome^10,12,107,108^. Grey fields indicate sequence matches using reciprocal TBLASTN between the predicted protein of a first species and the NCBI RefSeq_RNA dataset of the second species, but are only shown for flanking syntenic genes (green). Ser3 homologs (magenta) all show a high degree of similarity dominated by serine-rich repeats. **B-C.** Protein alignments of Ser3 orthologs from genome annotations of pyralid and crambid species (Pyraloidae), including copies of Ser3a previously dubbed as Ser3 in *E. kuehniella* and *A. transitella*^10^, with a focus on the N-terminal signal peptide (B) and a central repetitive region (C). Gene and protein identifiers are listed in **Table S7**. At : *Amyelois transitella* (Pyralidae, Phycitinae); Ek : *Ephestia kuehniella* (Pyralidae, Phycitinae); Pi : *Plodia interpunctella* (Pyralidae, Phycitinae); Gm : *Galleria mellonella* (Pyralidae, Galleriinae); Of : *Ostrinia furnacalis* (Crambidae, Pyraustinae).

**Figure S6.**
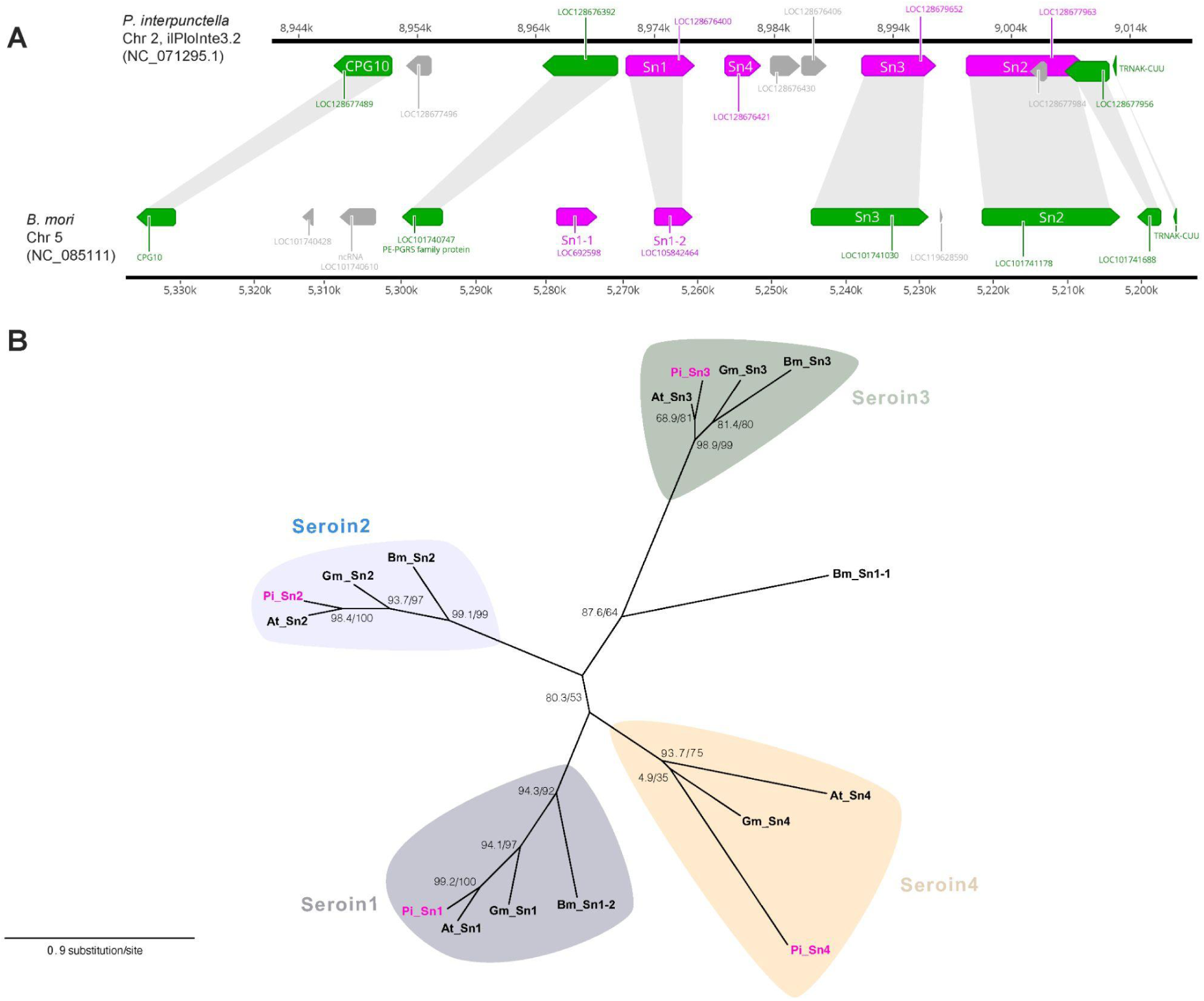
Homology and microsynteny of seroin genes in Lepidoptera. **A.** Synteny comparison between the clusters encompassing four seroin genes in *P. interpunctella* (top) and *B. mori* (bottom). Grey fields indicate sequence matches using reciprocal TBLASTN between the predicted protein of a first species and the NCBI RefSeq_RNA dataset of the second species. **B.** Maximum likelihood phylogenetic reconstruction of pyralid and *B. mori* seroin proteins, highlighting four orthology groups. Branch support is indicated by SH-aLRT % values / ultrafast bootstrap % values. Gene and protein identifiers are listed in **Table S7**. At : *Amyelois transitella* (Pyralidae, Phycitinae); Bm : *Bombyx mori* (Bombycidae, Bombycinae); Gm : *Galleria mellonella* (Pyralidae, Galleriinae) ; Pi : *Plodia interpunctella* (Pyralidae, Phycitinae).

**Table S1**: HCR Oligonucleotides Sequences

**Table S2** : DESeq2 Results Table

**Table S3** : Table containing the DESeq2 normalized counts for all annotated genes in the *ilPloInte3.2* genome

**Table S4** : Table containing data plotted in the Heatmap, related to Figure 3

**Table S5** : Table containing the TPM normalized counts for all annotated genes in the *ilPloInte3.2* genome

**Table S6** : Table containing data plotted in the Scatter Plot, related to Figure 3

**Table S7** : Accession numbers for sericin factor genes used in synteny analyses and alignments

**Table S8**: Amino acid and signal peptide sequences for top MSG and PSG enriched genes

**Table S9**: Manually Curated *ilPloInte3.2* Gene List

**File S1**: Manually Curated *ilPloInte3.2* Genome Annotation GTF

**Video S1**: 3D rendering of silk glands in a fifth instar moth larva visualized by micro-CT.. Related to Figure 1

**Video S2**: XY-plane micro-CT slices through a fifth instar moth larva. Animated sequence of consecutive 2D micro-CT sections scrolling through the larva from anterior to posterior. Slices highlight internal anatomical structures including the changes in diameter, looping, and crossing of the silk glands along the larva’s body axis. Related to Figure 1.

**Video S3**: 3D projection of DAPI, Phalloidin, and WGA stainings, from confocal microscopy stacks of the ASG, MSG, and PSG, related to Figure 1

## References

1. Sutherland, T.D., Young, J.H., Weisman, S., Hayashi, C.Y., and Merritt, D.J. (2010). Insect silk: one name, many materials. Annu Rev Entomol 55, 171–188. 10.1146/annurev-ento-112408-085401.

2. Craig, C.L. (1997). Evolution of arthropod silks. Annu Rev Entomol 42, 231–267. 10.1146/annurev.ento.42.1.231.

3. Yonemura, N., and Sehnal, F. (2006). The design of silk fiber composition in moths has been conserved for more than 150 million years. J Mol Evol 63, 42–53. 10.1007/s00239-005-0119-y.

4. Fedic, R., Zurovec, M., and Sehnal, F. (2002). The silk of Lepidoptera. Journal of Insect Biotechnology and Sericology 71, 1–15.

5. Gamo, T., Inokuchi, T., and Laufer, H. (1977). Polypeptides of fibroin and sericin secreted from the different sections of the silk gland in Bombyx mori. Insect Biochemistry 7, 285–295.

6. Kono, N., Nakamura, H., Tateishi, A., Numata, K., and Arakawa, K. (2021). The balance of crystalline and amorphous regions in the fibroin structure underpins the tensile strength of bagworm silk. Zoological Lett 7, 11. 10.1186/s40851-021-00179-7.

7. Aikman, E.L., Eccles, L.E., and Stoppel, W.L. (2025). Native Silk Fibers: Protein Sequence and Structure Influences on Thermal and Mechanical Properties. Biomacromolecules 26, 2043–2059. 10.1021/acs.biomac.4c01781.

8. Malay, A.D., Sato, R., Yazawa, K., Watanabe, H., Ifuku, N., Masunaga, H., Hikima, T., Guan, J., Mandal, B.B., Damrongsakkul, S., et al. (2016). Relationships between physical properties and sequence in silkworm silks. Sci Rep 6, 27573. 10.1038/srep27573.

9. Reizabal, A., Costa, C.M., Pérez-Álvarez, L., Vilas-Vilela, J.L., and Lanceros-Méndez, S. (2023). Silk Fibroin as Sustainable Advanced Material: Material Properties and Characteristics, Processing, and Applications. Adv Funct Materials 33, 2210764. 10.1002/adfm.202210764.

10. Wu, B.C.-H., Sauman, I., Maaroufi, H.O., Zaloudikova, A., Zurovcova, M., Kludkiewicz, B., Hradilova, M., and Zurovec, M. (2022). Characterization of silk genes in Ephestia kuehniella and Galleria mellonella revealed duplication of sericin genes and highly divergent sequences encoding fibroin heavy chains. Front Mol Biosci 9, 1023381. 10.3389/fmolb.2022.1023381.

11. Sparkes, J., and Holland, C. (2018). The rheological properties of native sericin. Acta biomaterialia 69, 234–242.

12. Kludkiewicz, B., Kucerova, L., Konikova, T., Strnad, H., Hradilova, M., Zaloudikova, A., Sehadova, H., Konik, P., Sehnal, F., and Zurovec, M. (2019). The expansion of genes encoding soluble silk components in the greater wax moth, Galleria mellonella. Insect Biochem Mol Biol 106, 28–38. 10.1016/j.ibmb.2018.11.003.

13. Zhu, L.J., Arai, M., and Hirabayashi, K. (1995). Relationship between adhesive properties and structure of sericin in cocoon filaments. The Journal of Sericultural Science of Japan 64, 420–426. 10.11416/kontyushigen1930.64.420.

14. Brookstein, O., Shimoni, E., Eliaz, D., Dezorella, N., Biran, I., Rechav, K., Sivan, E., Kozell, A., and Shimanovich, U. (2024). The Natural Material Evolution and Stage-wise Assembly of Silk Along the Silk Gland. bioRxiv, 2024–04.

15. Julien, E., Coulon-Bublex, M., Garel, A., Royer, C., Chavancy, G., Prudhomme, J.-C., and Couble, P. (2005). Silk Gland Development and Regulation of Silk Protein Genes. In Comprehensive Molecular Insect Science (Elsevier), pp. 369–384. 10.1016/B0-44-451924-6/00022-3.

16. Walker, A.A., Holland, C., and Sutherland, T.D. (2015). More than one way to spin a crystallite: multiple trajectories through liquid crystallinity to solid silk. Proc Biol Sci 282, 20150259. 10.1098/rspb.2015.0259.

17. Brough, H.D.A., Cheneler, D., and Hardy, J.G. (2024). Progress in Multiscale Modeling of Silk Materials. Biomacromolecules 25, 6987–7014. 10.1021/acs.biomac.4c01122.

18. Markee, A., Godfrey, R.K., Frandsen, P.B., Weng, Y.-M., Triant, D.A., and Kawahara, A.Y. (2024). De Novo Long-Read Genome Assembly and Annotation of the Luna Moth (Actias luna) Fully Resolves Repeat-Rich Silk Genes. Genome Biology and Evolution 16, evae148.

19. Kawahara, A.Y., Storer, C.G., Markee, A., Heckenhauer, J., Powell, A., Plotkin, D., Hotaling, S., Cleland, T.P., Dikow, R.B., Dikow, T., et al. (2022). Long-read HiFi sequencing correctly assembles repetitive heavy fibroin silk genes in new moth and caddisfly genomes. GigaByte 2022, gigabyte64. 10.46471/gigabyte.64.

20. Moreno-Tortolero, R.O., Luo, Y., Parmeggiani, F., Skaer, N., Walker, R., Serpell, L.C., Holland, C., and Davis, S.A. (2024). Molecular organization of fibroin heavy chain and mechanism of fibre formation in Bombyx mori. Commun Biol 7, 786. 10.1038/s42003-024-06474-1.

21. Mi, J., Zhou, Y., Ma, S., Zhou, X., Xu, S., Yang, Y., Sun, Y., Xia, Q., Zhu, H., Wang, S., et al. (2023). High-strength and ultra-tough whole spider silk fibers spun from transgenic silkworms. Matter 6, 3661–3683. 10.1016/j.matt.2023.08.013.

22. Tanaka, K., Kajiyama, N., Ishikura, K., Waga, S., Kikuchi, A., Ohtomo, K., Takagi, T., and Mizuno, S. (1999). Determination of the site of disulfide linkage between heavy and light chains of silk fibroin produced by Bombyx mori. Biochim Biophys Acta 1432, 92–103. 10.1016/s0167-4838(99)00088-6.

23. Zabelina, V., Takasu, Y., Sehadova, H., Yonemura, N., Nakajima, K., Sezutsu, H., Sery, M., Zurovec, M., Sehnal, F., and Tamura, T. (2021). Mutation in Bombyx mori fibrohexamerin (P25) gene causes reorganization of rough endoplasmic reticulum in posterior silk gland cells and alters morphology of fibroin secretory globules in the silk gland lumen. Insect Biochem Mol Biol 135, 103607. 10.1016/j.ibmb.2021.103607.

24. Chen, X., Wang, Y., Wang, Y., Li, Q., Liang, X., Wang, G., Li, J., Peng, R., Sima, Y., and Xu, S. (2022). Ectopic expression of sericin enables efficient production of ancient silk with structural changes in silkworm. Nat Commun 13, 6295. 10.1038/s41467-022-34128-5.

25. Chang, H., Cheng, T., Wu, Y., Hu, W., Long, R., Liu, C., Zhao, P., and Xia, Q. (2015). Transcriptomic analysis of the anterior silk gland in the domestic silkworm (Bombyxmori)–insight into the mechanism of silk formation and spinning. PLoS One 10, e0139424.

26. Dong, Z., Zhao, P., Zhang, Y., Song, Q., Zhang, X., Guo, P., Wang, D., and Xia, Q. (2016). Analysis of proteome dynamics inside the silk gland lumen of Bombyx mori. Sci Rep 6, 21158. 10.1038/srep21158.

27. Shi, R., Ma, S., He, T., Peng, J., Zhang, T., Chen, X., Wang, X., Chang, J., Xia, Q., and Zhao, P. (2019). Deep Insight into the Transcriptome of the Single Silk Gland of Bombyx mori. Int J Mol Sci 20, 2491. 10.3390/ijms20102491.

28. Kimoto, M., Tsubota, T., Uchino, K., Sezutsu, H., and Takiya, S. (2014). Hox transcription factor Antp regulates sericin-1 gene expression in the terminal differentiated silk gland of Bombyx mori. Dev Biol 386, 64–71. 10.1016/j.ydbio.2013.12.002.

29. Tsubota, T., Tomita, S., Uchino, K., Kimoto, M., Takiya, S., Kajiwara, H., Yamazaki, T., and Sezutsu, H. (2016). A Hox Gene, Antennapedia, Regulates Expression of Multiple Major Silk Protein Genes in the Silkworm Bombyx mori. J Biol Chem 291, 7087–7096. 10.1074/jbc.M115.699819.

30. Asakura, T., Umemura, K., Nakazawa, Y., Hirose, H., Higham, J., and Knight, D. (2007). Some observations on the structure and function of the spinning apparatus in the silkworm Bombyx mori. Biomacromolecules 8, 175–181. 10.1021/bm060874z.

31. Sehadova, H., Zavodska, R., Zurovec, M., and Sauman, I. (2021). The Filippi’s Glands of Giant Silk Moths: To Be or Not to Be? Insects 12, 1040. 10.3390/insects12111040.

32. Sehadova, H., Zavodska, R., Rouhova, L., Zurovec, M., and Sauman, I. (2021). The Role of Filippi’s Glands in the Silk Moths Cocoon Construction. Int J Mol Sci 22, 13523. 10.3390/ijms222413523.

33. Kono, N., Nakamura, H., Ohtoshi, R., Tomita, M., Numata, K., and Arakawa, K. (2019). The bagworm genome reveals a unique fibroin gene that provides high tensile strength. Commun Biol 2, 1–9. 10.1038/s42003-019-0412-8.

34. Dong, Z., Song, Q., Zhang, Y., Chen, S., Zhang, X., Zhao, P., and Xia, Q. (2016). Structure, evolution, and expression of antimicrobial silk proteins, seroins in Lepidoptera. Insect Biochem Mol Biol 75, 24–31. 10.1016/j.ibmb.2016.05.005.

35. Zhu, H., Zhang, X., Lu, M., Chen, H., Chen, S., Han, J., Zhang, Y., Zhao, P., and Dong, Z. (2020). Antibacterial mechanism of silkworm seroins. Polymers 12, 2985.

36. Kucerova, L., Zurovec, M., Kludkiewicz, B., Hradilova, M., Strnad, H., and Sehnal, F. (2019). Modular structure, sequence diversification and appropriate nomenclature of seroins produced in the silk glands of Lepidoptera. Sci Rep 9, 3797. 10.1038/s41598-019-40401-3.

37. Dong, Y., Dai, F., Ren, Y., Liu, H., Chen, L., Yang, P., Liu, Y., Li, X., Wang, W., and Xiang, H. (2015). Comparative transcriptome analyses on silk glands of six silkmoths imply the genetic basis of silk structure and coloration. BMC Genomics 16, 203. 10.1186/s12864-015-1420-9.

38. Su, H., Cheng, Y., Wang, Z., Li, Z., Stanley, D., and Yang, Y. (2015). Silk Gland Gene Expression during Larval-Pupal Transition in the Cotton Leaf Roller Sylepta derogata (Lepidoptera: Pyralidae). PLOS ONE 10, e0136868. 10.1371/journal.pone.0136868.

39. Tsubota, T., Yamamoto, K., Mita, K., and Sezutsu, H. (2016). Gene expression analysis in the larval silk gland of the eri silkworm Samia ricini. Insect Sci 23, 791–804. 10.1111/1744-7917.12251.

40. Tsubota, T., Yoshioka, T., Jouraku, A., Suzuki, T.K., Yonemura, N., Yukuhiro, K., Kameda, T., and Sezutsu, H. (2021). Transcriptomic analysis of the bagworm moth silk gland reveals a number of silk genes conserved within Lepidoptera. Insect Science 28, 885–900. 10.1111/1744-7917.12846.

41. Volenikova, A., Nguyen, P., Davey, P., Sehadova, H., Kludkiewicz, B., Koutecky, P., Walters, J.R., Roessingh, P., Provaznikova, I., Sery, M., et al. (2022). Genome sequence and silkomics of the spindle ermine moth, Yponomeuta cagnagella, representing the early diverging lineage of the ditrysian Lepidoptera. Commun Biol 5, 1281. 10.1038/s42003-022-04240-9.

42. Duan, J., Li, S., Zhang, Z., Yao, L., Yang, X., Ma, S., Duan, N., Wang, J., Zhu, X., and Zhao, P. (2023). A transcriptional atlas of the silk gland in Antheraea pernyi revealed by IsoSeq. Journal of Asia-Pacific Entomology 26, 102043.

43. Matsunami, K., Kokubo, H., Ohno, K., and Suzuki, Y. (1998). Expression pattern analysis of SGF-3/POU-M1 in relation to sericin-1 gene expression in the silk gland. Dev Growth Differ 40, 591–597. 10.1046/j.1440-169x.1998.t01-4-00003.x.

44. Dhawan, S., and Gopinathan, K.P. (2003). Expression profiling of homeobox genes in silk gland development in the mulberry silkworm Bombyx mori. Dev Genes Evol 213, 523–533. 10.1007/s00427-003-0357-1.

45. Parthasarathy, R., and Gopinathan, K.P. (2005). Comparative analysis of the development of the mandibular salivary glands and the labial silk glands in the mulberry silkworm, Bombyx mori. Gene Expr Patterns 5, 323–339. 10.1016/j.modgep.2004.10.006.

46. Ma, Y., Zeng, W., Ba, Y., Luo, Q., Ou, Y., Liu, R., Ma, J., Tang, Y., Hu, J., Wang, H., et al. (2022). A single-cell transcriptomic atlas characterizes the silk-producing organ in the silkworm. Nat Commun 13, 3316. 10.1038/s41467-022-31003-1.

47. Ma, Y., Li, Q., Tang, Y., Zhang, Z., Liu, R., Luo, Q., Wang, Y., Hu, J., Chen, Y., Li, Z., et al. (2024). The architecture of silk-secreting organs during the final larval stage of silkworms revealed by single-nucleus and spatial transcriptomics. Cell Reports 43. 10.1016/j.celrep.2024.114460.

48. Fang, S.-M., Hu, B.-L., Zhou, Q.-Z., Yu, Q.-Y., and Zhang, Z. (2015). Comparative analysis of the silk gland transcriptomes between the domestic and wild silkworms. BMC Genomics 16, 60. 10.1186/s12864-015-1287-9.

49. Eccles, L.E., Aikman, E.L., McTyer, J.B., Cruz, I.L.M., Richgels, A.L., and Stoppel, W.L. (2025). Exploring the functional properties of Plodia interpunctella silk fibers as a natural biopolymer for biomaterial applications. Materials Today Communications 42, 111416.

50. Shirk, B.D., Torres Pereira Meriade Duarte, I., McTyer, J.B., Eccles, L.E., Lateef, A.H., Shirk, P.D., and Stoppel, W.L. (2024). Harvesting Silk Fibers from *Plodia interpunctella:* Role of Environmental Rearing Conditions in Fiber Production and Properties. ACS Biomater. Sci. Eng. 10, 2088–2099. 10.1021/acsbiomaterials.3c01372.

51. Shirk, B.D., Heichel, D.L., Eccles, L.E., Rodgers, L.I., Lateef, A.H., Burke, K.A., and Stoppel, W.L. (2024). Modifying Naturally Occurring, Nonmammalian-Sourced Biopolymers for Biomedical Applications. ACS Biomater. Sci. Eng. 10, 5915–5938. 10.1021/acsbiomaterials.4c00689.

52. Tendolkar, A., Pomerantz, A.F., Heryanto, C., Shirk, P.D., Patel, N.H., and Martin, A. (2021). Ultrabithorax Is a Micromanager of Hindwing Identity in Butterflies and Moths. Front. Ecol. Evol. 9. 10.3389/fevo.2021.643661.

53. Heryanto, C., Hanly, J.J., Mazo-Vargas, A., Tendolkar, A., and Martin, A. (2022). Mapping and CRISPR homology-directed repair of a recessive white eye mutation in Plodia moths. iScience, 103885.

54. Heryanto, C., Mazo-Vargas, A., and Martin, A. (2022). Efficient hyperactive piggyBac transgenesis in Plodia pantry moths. Frontiers in Genome Editing 4, 57.

55. Shirk, B.D., Shirk, P.D., Furlong, R.B., Scully, E.D., Wu, K., and Siegfried, B.D. (2023). Gene editing of the ABC Transporter/White locus using CRISPR/Cas9-mediated mutagenesis in the Indian Meal Moth. J Insect Physiol 145, 104471. 10.1016/j.jinsphys.2022.104471.

56. Sorrell, E.L., and Lubkin, S.R. (2022). Bubble packing, eccentricity, and notochord development. Cells Dev 169, 203753. 10.1016/j.cdev.2021.203753.

57. Yang, H., Guo, Y., Wang, J., Tao, C., Cao, J., Cheng, T., and Liu, C. (2024). Bmgsb is involved in the determination of cell fate by affecting the cell cycle genes in the silk gland of Bombyx mori. Int J Biol Macromol 283, 136914. 10.1016/j.ijbiomac.2024.136914.

58. Hampoelz, B., Azou-Gros, Y., Fabre, R., Markova, O., Puech, P.-H., and Lecuit, T. (2011). Microtubule-induced nuclear envelope fluctuations control chromatin dynamics in Drosophila embryos. Development 138, 3377–3386. 10.1242/dev.065706.

59. Gage, L.P. (1974). Polyploidization of the silk gland of Bombyx mori. J Mol Biol 86, 97–108. 10.1016/s0022-2836(74)80010-0.

60. Henderson, S.C., and Locke, M. (1991). The development of branched silk gland nuclei. Tissue Cell 23, 867–880. 10.1016/0040-8166(91)90036-s.

61. Dhawan, S., and Gopinathan, K.P. (2003). Cell cycle events during the development of the silk glands in the mulberry silkworm Bombyx mori. Dev Genes Evol 213, 435–444. 10.1007/s00427-003-0343-7.

62. Buntrock, L., Marec, F., Krueger, S., and Traut, W. (2012). Organ growth without cell division: somatic polyploidy in a moth, Ephestia kuehniella. Genome 55, 755–763. 10.1139/g2012-060.

63. Prakash, A., Dion, E., Banerjee, T.D., and Monteiro, A. (2024). The molecular basis of scale development highlighted by a single-cell atlas of Bicyclus anynana butterfly pupal forewings. Cell Reports 43, 114147.

64. Loh, L.S., DeMarr, K.A., Tsimba, M., Heryanto, C., Berrio, A., Patel, N.H., Martin, A., McMillan, W.O., Wray, G.A., and Hanly, J.J. (2025). Lepidopteran scale cells derive from sensory organ precursors through a canonical lineage. Development 152, DEV204501. 10.1242/dev.204501.

65. Choi, H.M., Schwarzkopf, M., Fornace, M.E., Acharya, A., Artavanis, G., Stegmaier, J., Cunha, A., and Pierce, N.A. (2018). Third-generation in situ hybridization chain reaction: multiplexed, quantitative, sensitive, versatile, robust. Development 145, dev165753. 10.1242/dev.165753.

66. Durand, B., Drevet, J., and Couble, P. (1992). P25 gene regulation in Bombyx mori silk gland: two promoter-binding factors have distinct tissue and developmental specificities. Mol Cell Biol 12, 5768–5777. 10.1128/mcb.12.12.5768.

67. Takiya, S., Tsubota, T., and Kimoto, M. (2016). Regulation of Silk Genes by Hox and Homeodomain Proteins in the Terminal Differentiated Silk Gland of the Silkworm Bombyx mori. J Dev Biol 4, 19. 10.3390/jdb4020019.

68. Kimoto, M., Tsubota, T., Uchino, K., Sezutsu, H., and Takiya, S. (2015). LIM-homeodomain transcription factor Awh is a key component activating all three fibroin genes, fibH, fibL and fhx, in the silk gland of the silkworm, Bombyx mori. Insect Biochem Mol Biol 56, 29–35. 10.1016/j.ibmb.2014.11.003.

69. Wu, B.C.-H., Zabelina, V., Zurovcova, M., and Zurovec, M. (2024). Characterization and comparative analysis of sericin protein 150 in Bombyx mori. Sci Rep 14, 20990. 10.1038/s41598-024-71503-2.

70. Soneson, C., Love, M.I., and Robinson, M.D. (2015). Differential analyses for RNA-seq: transcript-level estimates improve gene-level inferences. F1000Res 4, 1521. 10.12688/f1000research.7563.2.

71. Cao, J., Zheng, H.-S., Zhang, R., Xu, Y.-P., Pan, H., Li, S., Liu, C., and Cheng, T.-C. (2023). Dimmed gene knockout shortens larval growth and reduces silk yield in the silkworm, Bombyx mori. Insect Mol Biol 32, 26–35. 10.1111/imb.12810.

72. Zhao, X.-M., Liu, C., Jiang, L.-J., Li, Q.-Y., Zhou, M.-T., Cheng, T.-C., Mita, K., and Xia, Q.-Y. (2015). A Juvenile Hormone Transcription Factor Bmdimm-Fibroin H Chain Pathway Is Involved in the Synthesis of Silk Protein in Silkworm, Bombyx mori. J Biol Chem 290, 972–986. 10.1074/jbc.M114.606921.

73. Yang, H., Xu, Y., Yuan, Y., Liu, X., Zhang, J., Li, J., Zhang, R., Cao, J., Cheng, T., and Liu, C. (2024). Identification and function of the Pax gene Bmgsb in the silk gland of Bombyx mori. Insect Mol Biol 33, 173–184. 10.1111/imb.12886.

74. Yang, H., Qin, X., Guo, Y., Tao, C., Cao, J., Cheng, T., and Liu, C. (2025). Bmgsb directly activates Bmubxn-4 to inhibit the DNA endoreplication and affect the cell fate in the silk gland of Bombyx mori. Int J Biol Macromol 308, 142335. 10.1016/j.ijbiomac.2025.142335.

75. Guo, K., Duan, J., Jing, X., Zhang, X., Ding, Q., Dong, Z., Xia, Q., and Zhao, P. (2025). Silk components and properties of the multilayer cocoon of the greater wax moth, Galleria mellonella. Insect Sci. 10.1111/1744-7917.70047.

76. Nishihara, S. (2020). Functional analysis of glycosylation using Drosophila melanogaster. Glycoconj J 37, 1–14. 10.1007/s10719-019-09892-0.

77. Ten Hagen, K.G., Tran, D.T., Gerken, T.A., Stein, D.S., and Zhang, Z. (2003). Functional characterization and expression analysis of members of the UDP-GalNAc:polypeptide N-acetylgalactosaminyltransferase family from Drosophila melanogaster. J Biol Chem 278, 35039–35048. 10.1074/jbc.M303836200.

78. Sinohara, H. (1979). Glycopeptides isolated from sericin of the silkworm, Bombyx mori. Comparative Biochemistry and Physiology Part B: Comparative Biochemistry 63, 87–91. 10.1016/0305-0491(79)90239-6.

79. Tran, D.T., Zhang, L., Zhang, Y., Tian, E., Earl, L.A., and Ten Hagen, K.G. (2012). Multiple members of the UDP-GalNAc: polypeptide N-acetylgalactosaminyltransferase family are essential for viability in Drosophila. J Biol Chem 287, 5243–5252. 10.1074/jbc.M111.306159.

80. Rouhova, L., Zurovcova, M., Hradilova, M., Sery, M., Sehadova, H., and Zurovec, M. (2024). Comprehensive analysis of silk proteins and gland compartments in Limnephilus lunatus, a case-making trichopteran. BMC Genomics 25, 472. 10.1186/s12864-024-10381-4.

81. Dong, Z., Xia, Q., and Zhao, P. (2023). Antimicrobial components in the cocoon silk of silkworm, Bombyx mori. Int J Biol Macromol 224, 68–78. 10.1016/j.ijbiomac.2022.10.103.

82. Stefanović, O., Vukajlović, F., Mladenović, T., Predojević, D., Čomić, L., and Pešić, S.B. (2020). Antimicrobial activity of Indian meal moth silk, Plodia interpunctella. Current Science 118, 1609–1614.

83. Vegliante, F., and Hasenfuss, I. (2012). Morphology and diversity of exocrine glands in lepidopteran larvae. Annu Rev Entomol 57, 187–204. 10.1146/annurev-ento-120710-100646.

84. Guo, P.-C., Dong, Z., Xiao, L., Li, T., Zhang, Y., He, H., Xia, Q., and Zhao, P. (2015). Silk gland-specific proteinase inhibitor serpin16 from the Bombyx mori shows cysteine proteinase inhibitory activity. Biochem Biophys Res Commun 457, 31–36. 10.1016/j.bbrc.2014.12.056.

85. Kodrík, D., Kludkiewicz, B., Navrátil, O., Skoková Habuštová, O., Horáčková, V., Svobodová, Z., Vinokurov, K.S., and Sehnal, F. (2013). Protease inhibitor from insect silk-activities of derivatives expressed in vitro and in transgenic potato. Appl Biochem Biotechnol 171, 209–224. 10.1007/s12010-013-0325-9.

86. Guo, K., Zhang, X., Zhao, D., Qin, L., Jiang, W., Hu, W., Liu, X., Xia, Q., Dong, Z., and Zhao, P. (2022). Identification and characterization of sericin5 reveals non-cocoon silk sericin components with high β-sheet content and adhesive strength. Acta Biomater 150, 96–110. 10.1016/j.actbio.2022.07.021.

87. Takasu, Y., Hata, T., Uchino, K., and Zhang, Q. (2010). Identification of Ser2 proteins as major sericin components in the non-cocoon silk of Bombyx mori. Insect Biochem Mol Biol 40, 339–344. 10.1016/j.ibmb.2010.02.010.

88. Alqassar, J.D., and Martin, A. (2025). Methods for the study of gene expression in the silk glands of the pantry moth Plodia interpunctella. 10.17605/OSF.IO/3PZ6C.

89. Bruce, H.S., Jerz, G., Kelly, S., McCarthy, J., Pomerantz, A., Senevirathne, G., Sherrard, A., Sun, D.A., Wolff, C., and Patel, N.H. (2021). Hybridization chain reaction (HCR) in situ protocol. protocols. io. 10.17504/protocols.io.bunznvf6.

90. Kuehn, E., Clausen, D.S., Null, R.W., Metzger, B.M., Willis, A.D., and Özpolat, B.D. (2022). Segment number threshold determines juvenile onset of germline cluster expansion in Platynereis dumerilii. Journal of Experimental Zoology Part B: Molecular and Developmental Evolution 338, 225–240. 10.1002/jez.b.23100.

91. Schindelin, J., Arganda-Carreras, I., Frise, E., Kaynig, V., Longair, M., Pietzsch, T., Preibisch, S., Rueden, C., Saalfeld, S., and Schmid, B. (2012). Fiji: an open-source platform for biological-image analysis. Nature methods 9, 676.

92. Chiu, C.-L., and Clack, N. (2022). Napari: a Python multi-dimensional image viewer platform for the research community. Microscopy and Microanalysis 28, 1576–1577.

93. Andrews, S. (2025). s-andrews/FastQC.

94. Chen, S., Zhou, Y., Chen, Y., and Gu, J. (2018). fastp: an ultra-fast all-in-one FASTQ preprocessor. Bioinformatics 34, i884–i890. 10.1093/bioinformatics/bty560.

95. Dobin, A., and Gingeras, T.R. (2016). Optimizing RNA-Seq mapping with STAR. In Data mining techniques for the life sciences (Springer), pp. 245–262.

96. Öztürk-Çolak, A., Marygold, S.J., Antonazzo, G., Attrill, H., Goutte-Gattat, D., Jenkins, V.K., Matthews, B.B., Millburn, G., dos Santos, G., Tabone, C.J., et al. (2024). FlyBase: updates to the Drosophila genes and genomes database. Genetics 227, iyad211. 10.1093/genetics/iyad211.

97. Liao, Y., Smyth, G.K., and Shi, W. (2014). featureCounts: an efficient general purpose program for assigning sequence reads to genomic features. Bioinformatics 30, 923–930. 10.1093/bioinformatics/btt656.

98. Darling, A.E., Mau, B., and Perna, N.T. (2010). progressiveMauve: multiple genome alignment with gene gain, loss and rearrangement. PLoS One 5, e11147. 10.1371/journal.pone.0011147.

99. Katoh, K., and Standley, D.M. (2013). MAFFT multiple sequence alignment software version 7: improvements in performance and usability. Molecular biology and evolution 30, 772–780.

100. Trifinopoulos, J., Nguyen, L.-T., von Haeseler, A., and Minh, B.Q. (2016). W-IQ-TREE: a fast online phylogenetic tool for maximum likelihood analysis. Nucleic acids research 44, W232–W235.

101. Love, M.I., Huber, W., and Anders, S. (2014). Moderated estimation of fold change and dispersion for RNA-seq data with DESeq2. Genome biology 15, 1–21.

102. Wickham, H., and Sievert, C. (2009). ggplot2: elegant graphics for data analysis (springer New York).

103. Vries, A. de, and Ripley, B.D. (2024). ggdendro: Create Dendrograms and Tree Diagrams Using “ggplot2.”

104. Teufel, F., Almagro Armenteros, J.J., Johansen, A.R., Gíslason, M.H., Pihl, S.I., Tsirigos, K.D., Winther, O., Brunak, S., von Heijne, G., and Nielsen, H. (2022). SignalP 6.0 predicts all five types of signal peptides using protein language models. Nat Biotechnol 40, 1023–1025. 10.1038/s41587-021-01156-3.

105. Rindos, M., Kucerova, L., Rouhova, L., Sehadova, H., Sery, M., Hradilova, M., Konik, P., and Zurovec, M. (2021). Comparison of Silks from Pseudoips prasinana and Bombyx mori Shows Molecular Convergence in Fibroin Heavy Chains but Large Differences in Other Silk Components. Int J Mol Sci 22, 8246. 10.3390/ijms22158246.

106. Rouhova, L., Kludkiewicz, B., Sehadova, H., Sery, M., Kucerova, L., Konik, P., and Zurovec, M. (2021). Silk of the common clothes moth, Tineola bisselliella, a cosmopolitan pest belonging to the basal ditrysian moth line. Insect Biochem Mol Biol 130, 103527. 10.1016/j.ibmb.2021.103527.

107. Zurovec, M., Kludkiewicz, B., Fedic, R., Sulitkova, J., Mach, V., Kucerova, L., and Sehnal, F. (2013). Functional conservation and structural diversification of silk sericins in two moth species. Biomacromolecules 14, 1859–1866. 10.1021/bm400249b.

108. Zurovec, M., Yang, C., Kodrík, D., and Sehnal, F. (1998). Identification of a novel type of silk protein and regulation of its expression. J Biol Chem 273, 15423–15428. 10.1074/jbc.273.25.15423.

109. Victoriano, E., and Gregório, E.A. (2004). Ultrastructure of the Lyonet’s glands in larvae of Diatraea saccharalis Fabricius (Lepidoptera: Pyralidae). Biocell 28, 165–169.

